# Modular cloning of multigene vectors for the baculovirus system and yeast

**DOI:** 10.1101/2024.10.07.616927

**Authors:** Zhihao Lai, Sarena F. Flanigan, Marion Boudes, Chen Davidovich

## Abstract

Recombinant macromolecular complexes are often produced by the baculovirus system, using multigene expression vectors. Yet, the construction of baculovirus-compatible multigene expression vectors is complicated and time-consuming. Furthermore, while the baculovirus and yeast are popular protein expression systems, no single method for multigene vector construction is compatible with both. Here we present the modular cloning (MoClo) Baculo toolkit for constructing multigene expression vectors for the baculovirus system and, through compatibility with the MoClo Yeast toolkit, also for yeast. Vector construction by MoClo Baculo utilises Golden Gate assembly, which does not require PCR, primers or the sequencing of intermediate products. As a proof of principle, MoClo Baculo was used to construct baculovirus and yeast multigene vectors expressing the four- and five-subunit human Polycomb Repressive Complex 2. We show that MoClo Baculo simplifies and expedites the construction of multigene expression vectors for the baculovirus system and provides compatibility with yeast as an alternative expression system.

## Introduction

An increasing number of large multi-subunit macromolecular complexes are subjected to structure determination using cryo-electron microscopy [1]. Such structural and biophysical studies often require the expression and purification of recombinant multisubunit protein complexes. When the macromolecule complex of interest is of a eukaryotic origin, its subunits are often required to be co-expressed in a eukaryotic host [2,3]. Among the most commonly used eukaryotic systems for recombinant protein expression are the baculovirus system [4] and yeast [5], each offering different advantages.

There are several widely used baculovirus expression systems. The Bac-to-Bac system is one of the most commonly used systems for the expression of protein complexes with a few protein subunits [6]. The Bac-to-Bac system allows the cloning of up to two genes per expression vector, which is then integrated into the baculovirus DNA and packed into baculovirus particles [6]. Complexes with more than two protein subunits can be expressed by the Bac-to-Bac system. This is commonly done by the co-infection of insect cells using multiple different baculoviruses, each carrying a different gene. Accordingly, there are many examples of protein complexes that were expressed using co-infection with 3-5 baculoviruses (e.g. [7,8,9,10,11]). Yet, beyond that, the efficiency and reproducibility of co-infections is reduced and, therefore, the expression of complexes with more than a few subunits commonly requires the usage of multi-gene vectors.

There are several multigene vector systems for protein expression using the baculovirus system. MultiBac recombines multiple genes into one vector using the Cre-lox technology [12]. BiGBac enables the construction of similar multigenic vectors using a Gibson Assembly [13], allowing a maximum of 25 genes in one plasmid [14]. GoldenBac utilises Golden Gate assembly that simplifies vector construction, where individual genes are PCR amplified only once, at the beginning of the process [15].

All current multigene baculovirus expression systems [6,11,12,14,15] require PCR or Gibson Assembly [13], and therefore oligonucleotide synthesis. PCR-based approaches may complicate the usage of GC-rich or repetitive templates for the construction of some multigene vectors. Furthermore, most multigene baculovirus vectors do not have a protein tag pre-incorporated in the vector. Instead, large protein tags are often added through a prior cloning step while small tags are included in the cloning primers. These factors increase the reliance on oligonucleotide synthesis and the subsequent sequencing of intermediate products, which increase the time, cost and complexity of the project.

The problem of multigene vector construction has already been solved in various organisms using modular cloning (MoClo) [16]. In MoClo, different ‘parts’ are functional DNA sequence elements that are assembled together to form ‘devices’, which are more complex plasmids. Each ‘part’ is a plasmid that includes one functional element, such as a protein-coding sequence, tag, promoter, or backbone of an expression vector. Each part is generated only once, usually with the aid of gene synthesis or gBlocks, and can then be used repeatedly in different projects or shared between labs. Multiple plasmids, each holding a single part, are digested together by type IIS restriction enzymes to release DNA fragments with cohesive ends that allow multiple parts to assemble together through Golden Gate assembly [17].

Golden Gate assembly uses Type IIS restriction enzymes, which cut outside of the enzyme recognition site. Thus, one enzyme can produce many different cohesive ends within the same reaction, while only compatible cohesive ends adhere and ligate to each other within a single-pot reaction. Accordingly, Golden Gate assembly can be used for the digestion of multiple plasmids or DNA fragments and their simultaneous assembly into a single vector. The restriction sites are destroyed upon ligation and, therefore, the product cannot be digested once ligated, making the Golden Gate assembly highly efficient as it typically progresses in one direction.

MoClo approaches are commonly based on multiple Golden Gate assembly reactions that are carried out iteratively. For instance, multiple plasmids can be assembled in parallel, using multiple independent Golden Gate assemblies with the aid of the same type IIS restriction enzyme. Subsequently, the obtained plasmids can be combined in another Golden Gate assembly, using another type IIS enzyme, where they are ligated again to form an increasingly complex plasmid that includes multiple functional elements. MoClo-constructed plasmids may include multiple genes and all the elements that are required for their delivery, integration into the host, selection, protein expression and protein purification. MoClo is fast, efficient, affordable, and does not require PCR, Gibson Assembly or custom primer design.

The MoClo Yeast Toolkit [18] was designed for modular cloning of multigene expression cassettes, aiming for synthetic biology applications in the yeast *S. cerevisiae*. Although the MoClo Yeast Toolkit was not designed primarily for the expression and purification of recombinant proteins, we reasoned that it would be feasible to modify it for that purpose. Based on a cloning standard inspired by the MoClo Yeast Toolkit, we set out to develop a method for cross-platform multi-gene vector building for protein expression and purification. We focused on constructing parts for modular cloning of expression vectors dedicated to the baculovirus system and yeast, given their broad applicability.

Here we present the MoClo Baculo, a multi-gene modular cloning (MoClo) method designed for the expression of proteins using the baculovirus system and in yeast. MoClo Baculo utilises Golden Gate as the only cloning method. The mutation rate is extremely low, given that MoClo does not involve the usage of PCR, Gibson Assembly, *in vitro* polymerases or primers. Therefore, the sequencing of intermediate product plasmids is not required, which dramatically reduces the time, cost and complexity of building multigene constructs. Modular cloning allows for flexible replacement or removal of N- or C-terminal affinity tags when developing a new purification scheme. MoClo Baculo supports the expression of up to 6 proteins from a single multigene construct, however this number can theoretically be expanded by adding adapters. The number of co-expressed subunits can further be expanded through the delivery of multiple multigene constructs. As a proof of concept, we used MoClo Baculo to build multi-gene expression vectors of the 4- and 5-subunits Polycomb repressive complex 2 (PRC2). We demonstrate the suitability of such MoClo Baculo expression vectors to express catalytically active PRC2 complexes using the baculovirus system and yeast.

## Results

### Modification of the MoClo Yeast toolkit for better compatibility with applications of recombinant protein expression and purification

We set out to generate a system that would utilise the same set of open reading frames (ORFs) and tag constructs for building different multi-protein expression vectors. Furthermore, we wished to have compatibility with protein expression using the baculovirus system and yeast. We prioritised methods that do not rely on PCR, in vitro polymerase reactions or primers, aiming to avoid the sequencing of intermediate products. We reasoned that the MoClo Yeast Toolkit [18], originally developed for synthetic biology applications, would be an ideal starting point.

The MoClo Baculo toolkit design is inherited from the MoClo Yeast Toolkit and therefore includes the following part types that are assembled together to construct expression vectors (Supplementary Fig. 1 and Supplementary Table 1): Left Connector (ConL; part 1) that connects functional elements to the plasmid backbone; Promoter, designed for protein expression with (part 2a) or without (part 2) N-terminal tag; an optional N-terminal tag (part 2b); a protein Coding Sequence (CDS; part 3); an optional C-terminal tag (part 4a); a transcription Terminator sequence that is designed to be used with or without C-terminal tag (part 4b or 4, respectively); a right Connector (ConR; part 5) and a backbone (part 6) that can be divided into up to 4 different parts (parts 6a-d) with different functionalities (more below).

One important difference between the MoClo Yeast Toolkit and the MoClo Baculo designs is that in the latter one can use the same CDS part with or without tags (Supplementary Fig. 1 and Supplementary Table 1). Specifically, in the MoClo Yeast toolkit, the construction of vectors for expressing untagged and N-terminally tagged proteins is done using two different CDS parts (Supplementary Fig. 1). This design is undesirable for structural biologists, who would benefit from the frequent removal or inclusion of N-terminal tags without having to source two separate CDS parts. Therefore, to allow for removing the N-terminal tag without changing the CDS part, we modified the MoClo Yeast general design of promoters and N-terminal parts (Supplementary Fig. 1A).

To enable the modular cloning using Golden Gate assembly, each part is domesticated into a chloramphenicol resistance plasmid (pYTK001) that originated from the MoClo Yeast toolkit (Supplementary Fig. 1D). The resulting plasmid holds a single part and is referred to as “Level 0” plasmid [18] (see methods section and Supplementary Fig. 1C,D for the flanking sequences that are added to each part, to allow for its domestication and subsequent usage).

The newly modified design of MoClo Baculo still allows the replacement of promoters through modular cloning, although it is not fully compatible with the promoter design of the MoClo Yeast toolkit [18]. Therefore, we generated several useful MoClo Baculo-compatible *S. cerevisiae* promoters, based on promoter sequences from the MoClo Yeast toolkit (Supplementary Table 1). More *S. cerevisiae* promoters can be made as needed. We also designed and generated parts carrying N- and C-terminal tags that are useful for protein expression and purification. Such tags for protein purification include a PreScission (3C) protease cleavable maltose binding protein (MBP) tag, a dual protein A tag and a Strep-tag (Supplementary Table 1). Additional tags can readily be added as individual parts, without bringing the coding sequence into consideration and as long as the correct flanking sequences are added to them (Supplementary Fig. 1C,D).

### Adding compatibility with the baculovirus system for protein expression

We next sought to enable compatibility between the MoClo Yeast toolkit to the baculovirus system. To this end, we modified parts of an expression vector used in the Bac-to-Bac expression system (pFastBac1 [6]), aiming to enable modular cloning. For that, we first aligned backbone sequences from different existing baculovirus expression systems that were previously modified from the Bac-to-Bac systems. This was done to identify commonly used modules (Supplementary Fig. 1B) and regions where the pFastBac1 vector can be modified without negatively impacting performances. Based on this sequence analysis we designed two new parts that originated from the pFastBac1 vector and are termed here baculovirus (BV) ‘Backbone’ (BV BB in Fig. 1, part 6b-d in Supplementary Table 1 and plasmid pMB.001 in Supplementary Table 2) and ‘Recombination into BV DNA’ (BV rec. in Fig. 1, part 6a in Supplementary Table 1, and plasmid pF003 in Supplementary Table 2). The BV Backbone (BV BB, part 6b-d; plasmid pMB.001) includes bacterial replication origin, ampicillin and gentamicin resistance and the right transposition site (Tn7R), which is used for recombination into the baculovirus DNA. The left transposition site (Tn7L) is included within the ‘Recombination into BV DNA’ (BV rec.) part (part 6a; plasmid pF003). Together, the ‘Backbone’ (BV BB) part and the ‘Recombination into BV DNA’ (BV rec.) part include all the functional elements required to connect to each other and to all the other parts that are collectively required to express a single open reading frame, either tagged or not. We envision that the usage of two parts for the backbone would facilitate further backbone modifications in future works. Details regarding adding new parts can be found in Supplementary Table 3. For compatibility with the MoClo Yeast cloning standard, we used codon optimization to remove BsaI and BsmBI sites from the sequences that were used to construct all these parts. In the case of parts that are specific for yeast, or may be used for both yeast and baculovirus expression, we also removed NotI sites, as NotI is used to linearise vectors for yeast genome integration. If one wishes to use the optional compatibility between MoClo Baculo to the biGBac system (more below), then PmeI sites should be removed too.

**Figure 1.**
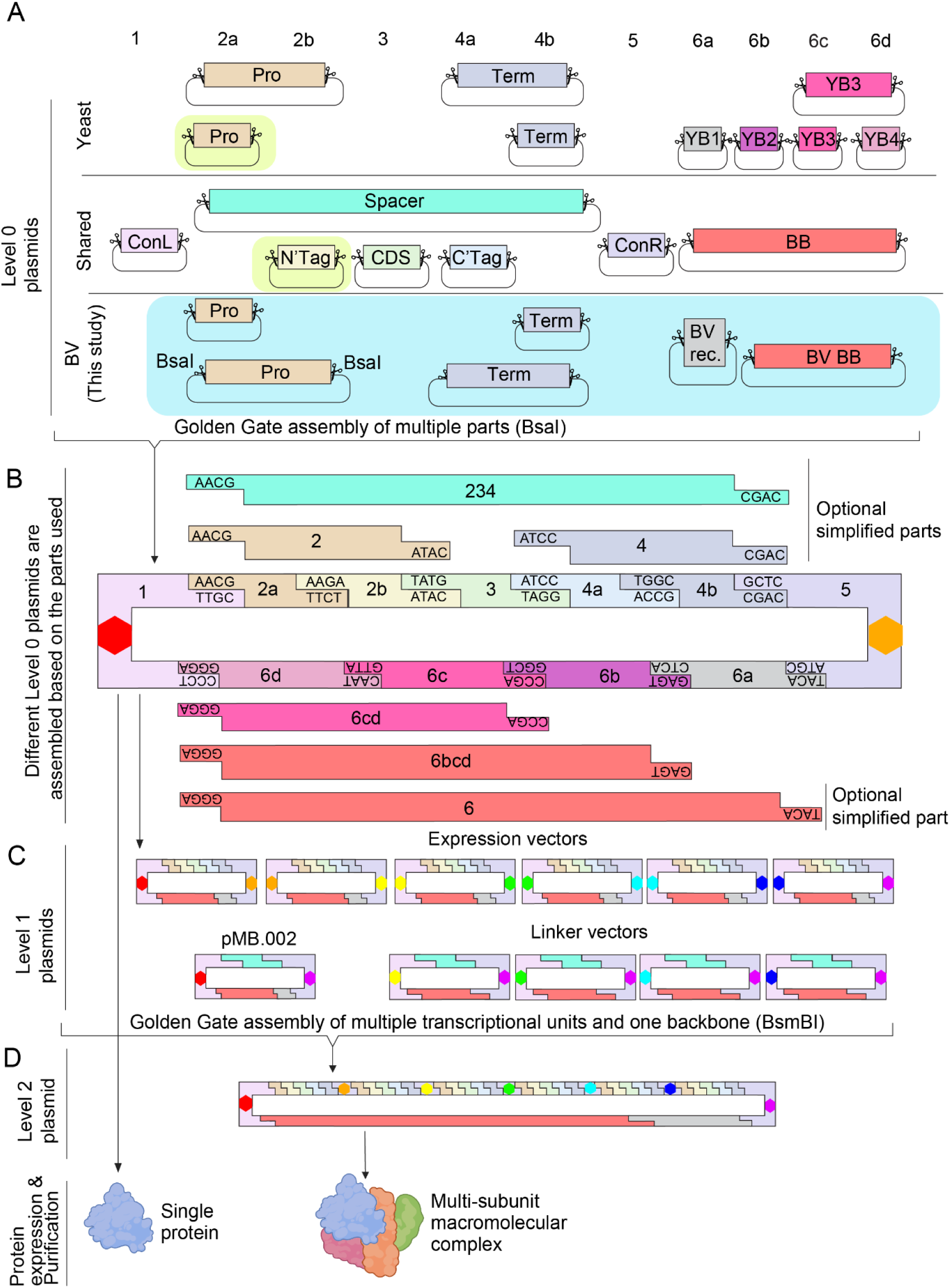
Design of the MoClo Baculo system. (A) Level 0 plasmids carrying individual parts, each with a different function as indicated. ConL and ConR: Connectors designed to connect the left (ConL) and right (ConR) assemblies of parts originated from multiple Level 1 plasmids. Pro: a promoter compatible with the intended host organism. Promoters are available as type 2a or 2 parts, used in the presence or absence of an N-terminal tag, respectively. N’ Tag: optional N-terminal tag. CDS: coding sequence of the protein of interest. C’tag: optional C-terminal tag. Term: transcription termination sequence. The terminator is available as type 4b or 4 part, used in the presence or absence of a C-terminal tag, respectively. YB1: yeast antibiotic resistance. YB2: 3’ homologous arm, aiming for chromosomal integration in yeast, or for yeast plasmid maintenance. YB3: origin of replication (ori) for bacterial plasmid together with bacterial antibiotic selection marker. YB4: 5’ homologous arm, aiming for chromosomal integration in yeast. BB: origin of replication for bacterial plasmid together with bacterial antibiotic selection marker. BV rev.: Tn7L for bacmid recombination. BV BB: all elements from BB and also gentamicin resistance (GentR) and Tn7R for bacmid recombination. Parts created in this study for BV expression are highlighted in light blue. Parts created in this study for compatibility between the MoClo Baculo to the MoClo Yeast Toolkits were highlighted in light green (see Supplementary Fig. 1A). For the assembly of Level 1 plasmids, parts are subjected to Golden Gate assembly using the BsaI restriction enzyme (scissors). (B) The level 0 plasmids are cleaved using BsaI into fragments with complementary overhangs, as indicated, and these DNA fragments are ligated using a Golden Gate assembly. Level 1 plasmids are transcription units, each capable of expressing a single protein. An exception is where a simple backbone (BB, part 6) is used for the assembly of the transcription unit, which leads to a transcription unit that can be assembled into a functional Level 2 expression vector, although the Level 1 plasmid is not designed to function as an expression vector by itself. The same colour scheme as in A was used. Sites that are dedicated to restriction digestion using BsmBI, during the subsequent Golden Gate assembly of Level 2 plasmids, were marked as colourful hexagons, with the type of overhang coded by its colour (see illustration in the subsequent panel). (C) Multiple Level 1 vectors, including multiple transcription units and one backbone, are subjected to Golden Gate assembly using the BsmBI enzyme to form one Level 2 vector (only bases that are marked by same-colour hexagons can be ligated, using cohesive ends that are formed upon BsmBI cleavage). In some cases, a ‘Linker’ vector can replace a transcription unit in order to link between another transcription unit to the backbone (bottom right in (C)). (D) Level 2 plasmids are multi-gene expression vectors, designed to express a multi-subunit macromolecular complex. The compatible expression system is determined based on the parts selected in (A). Yeast expression vectors were designed to express proteins either after homologous recombination into a chromosome or using a yeast plasmid. The order of the transcription units is determined based on the connectors that were selected in (A).

For compatibility with the baculovirus expression system, we designed additional new parts using the MoClo cloning standard: polyhedrin promoter (parts 2 and 2a) and SV40 terminator (parts 4 and 4b). All these parts were domesticated into Level 0 plasmids and are compatible with Golden Gate assembly using the type IIS restriction enzyme BsaI. At this point, the system included all the elements required to assemble expression vectors that are compatible with the Bac-to-Bac system. Each of the resulting expression vectors are termed “Level 1” plasmids (Fig. 1, top and Supplementary Fig. 1A). Level 1 plasmids can express a single protein from an element that is termed a “Transcription Unit”: a functional element required to transcribe an mRNA that is coding for the expression of a desired protein.

To enable the assembly of multiple Transcription Units (from multiple Level 1 plasmids) into a single multi-gene baculovirus expression vector (termed “Level 2” plasmid), we required a new backbone that confers additional antibiotic resistance. For that, we added spectinomycin resistance into the ‘Recombination into BV DNA’ part (BV rec., part 6a). The new part 6a (‘Recombination into BV DNA + SpecR’ [BV rec. + SpecR]) can now be assembled together with the Backbone (BV BB, part 6b-d), a Spacer (part 2-4) and Connectors (parts 1 and 5) through a Golden Gate reaction using BsaI. The result is a Level 1 plasmid that is termed here ‘Level 2 Backbone’. The Level 2 Backbone plasmid does not carry a Transcription Unit but, instead, contributes the backbone during the assembly of Level 2 plasmids. Users can assemble their own ‘Level 2 Backbone’ plasmids using parts from the MoClo Baculo toolkit, together with Connector parts from the MoClo Yeast toolkit. However, for the ease of use, we included in the MoClo Baculo toolkit a Level 2 Backbone plasmid (pMB.002) that includes the ConLS’ and ConRE’ connectors. The pMB.002 plasmid allows the integration of up to 6 transcription units into a Level 2 plasmid (Fig. 1B, right, and Supplementary Table 2). Overall, this design ensures that both single-gene (Level 1) and multi-gene (Level 2) expression vectors are suitable for protein expression using the Bac-to-Bac system and, therefore, confer resistance to ampicillin and gentamicin. In addition to these, multi-gene (Level 2) baculovirus expression vectors also confer resistance to spectinomycin.

### Modular cloning of baculovirus expression vectors

From this point, modular cloning is carried out using the same workflow that has been described for the MoClo Yeast toolkit [18]. As the process is described below in brief, it is highly recommended to read the description of modular cloning by Weber et al [16] and Lee et al [18]. Briefly, all the Level 0 (chloramphenicol resistant) vectors that carry the parts for a desired Transcription Unit are combined in a single pot Golden Gate assembly, using the type IIS restriction enzyme BsaI (Fig 1A). The produced Level 1 plasmid has lost its chloramphenicol resistance and now both the yeast and baculovirus expression vectors confer resistance to ampicillin in *E. coli*. The Level 1 baculovirus expression vectors also confer resistance to gentamicin in *E. coli*. This transition between antibiotic selection markers is inherited from the modular cloning method and allows selection against precursor plasmids while progressing up to the next plasmid Level through iterative cloning. For the production of baculovirus expression vectors, we use chloramphenicol to select Level 0 plasmids, ampicillin or gentamicin to select Level 1 plasmids and spectinomycin to select Level 2 plasmids. Each Level 1 plasmid can now be used to express a single protein, depending on the transcription unit and the backbone. Multiple Level 1 plasmids can be used to assemble one Level 2 plasmid—a multi-gene expression vector—using the Golden Gate assembly region with BsmBI digestion (Fig. 1B, bottom). Yet, the assembly of Level 2 vectors requires some planning, because the order of the transcription units in a Level 2 vector is defined based on the Connectors (parts 1 and 5) that are included in each of the Level 1 vectors that are combined to assemble a multi-gene expression vector (Level 2).

Lee et al. [18] and Weber et al. [16] are highly recommended readings on the topic of connector design, as the MoClo Baculo toolkit uses the connector parts of the MoClo Yeast toolkit [18]. In brief, connectors are designed such that a right Connector (ConR, part 5) of a given Transcription Unit can be connected only to a complementary left Connector (ConL, part 1) of another Transcription unit. Exceptions are the ConLS’ and ConRE’ connectors that are reserved for the backbone. For instance, ConR1 is connected to ConL1 of another Transcription Unit, ConR2 is connected to ConL2 in another Transcription unit, and so forth (see Supplementary Table 1 for the list of connectors). Practically, this means that the order of the transcription units in a given Level 2 vector usually need to be defined at the time when one designs the relevant set of Level 1 vectors that are intended to be used together in a subsequent Golden Gate assembly.

Even when Level 2 expression vector is intended for the baculovirus system, we find it convenient to use the same connector parts that were designed by Lee et al for the MoClo Yeast toolkit [18]. The MoClo Yeast toolkit provides connectors for combining up to 6 transcription units together. If less than 6 subunits are assembled, a linker plasmid can be used to connect the last transcription unit to the backbone (an example for that is described in the next section). It is anticipated that more connectors can be added to increase the number of transcription units in a given multi-gene expression vector, as long as the total length of the construct permits Golden Gate assembly.

We find that the sequencing of Level 1 plasmids is not required if they are only used as intermediate products for assembling a Level 2 vector. This is because mutations are uncommon, since no PCR or oligonucleotides are used. Therefore, when Level 1 plasmids are only used as intermediate products, we commonly validate the correct Level 1 plasmid assembly by restriction digestion before proceeding to Level 2 assembly. At the end of the process, the correct assembly of the Level 2 plasmid is validated by restriction digestion followed by whole plasmid sequencing. The obtained Level 2 plasmids are multi-gene expression vectors compatible with the bac-to-bac baculovirus expression system (Fig. 1C).

### Adding compatibility with the biGBac system

We envisioned that some users may benefit from compatibility between MoClo Baculo and the biGBac system [14]. This is because the biGBac system allows to connect together 5 multi-gene vectors using Gibson Assembly. Therefore, we decided to generate MoClo Baculo parts that would include the Gibson Assembly linkers of the biGBac system. For that, we modified the pMB.002 plasmid by inserting linker sequences (“A-F”) from pBIG1a-e [14] and PmeI restriction enzyme sites (Supplementary Fig. 2A). We named the resulting plasmids pMB.BIG1a to pMB.BIG1e. These plasmids are functionally identical to the biGBac system plasmids pBIG1a to pBIG1e plasmids, respectively. The compatibility between MoClo Baculo to the biGBac system may allow the optional connection of up to five Level 2 MoClo Baculo plasmids, each with up to six transcription units. The resulting biGBac expression vectors are referred to as Level 3 plasmids here and contain up to 30 transcription units.

We believe that most users will not need the compatibility with the biGBac system, and therefore it is marked here as optional. This is because compatibility with biGBac is beneficial only if one wishes to increase the maximal number of transcriptional units from six (Level 2 plasmids) to 30 (optional Level 3; Supplementary Fig. 2B) without adding new MoClo Baculo adapters. However, the added compatibility between the MoClo Baculo to the biGBac system may allow regular users of the biGBac system to construct their expression vectors with the aid of Modular Cloning. This may provide biGBac users with the simplicity of removing or replacing tags without PCR or gibson assembly.

### Modular cloning of yeast expression vectors

The construction of yeast expression vectors is carried out using the same CDS and tag parts that are used for baculovirus vector building. The process of vector building for yeast expression is nearly identical to the construction of baculovirus vectors, as described above. An exception is that for the construction of yeast expression vectors we normally use chloramphenicol to select Level 0 plasmids, ampicillin to select Level 1 plasmids and kanamycin to select Level 2 plasmids, as described by Lee et al [18]. Most of the parts that are used for building Level 1 and 2 plasmids for yeast expression are taken from the MoClo Yeast toolkit [18] (Fig. 1A; boxes over a white background). Exceptions are the following parts that were generated for the MoClo Baculo toolkit (Fig. 1A; boxes over a green or blue background): (i) Yeast promoter parts dedicated for using together with N-terminal tags according to the MoClo Baculo cloning standard and (ii) the same N-terminal tag parts; (iii) CDS parts and (iv) C-terminal tag parts that are both used for baculovirus expression but are also compatible with the yeast expression vectors.

### Proof of principle: multi-gene vector construction

As MoClo Baculo adapted the Modular Cloning strategy of the MoClo Yeast toolkit [18], the efficiency of cloning Level 0 and Level 1 vectors is close to 100 %. As a demonstration of Level 1 cloning, we constructed six different Level 1 plasmids (Fig. 2A). These included a backbone plasmid for baculovirus expression (pMB.002) and five Transcription Units for the expression of each of the five subunits of the PRC2-AEBP2 complex (Fig. 2A,B). These five Transcription Units designed for the expression of the PRC2 protein subunits MBP-EZH2, MBP-SUZ12, MBP-EED MBP-RBBP4 and AEBP2, where MBP is a PreScission-cleavable maltose binding protein tag. After the Golden Gate assembly reaction, two clones were picked per each of these Level 1 plasmids. All these clones were then assayed using restriction digestion, confirming a success rate of 100% (12/12; Fig 2A).

**Figure 2.**
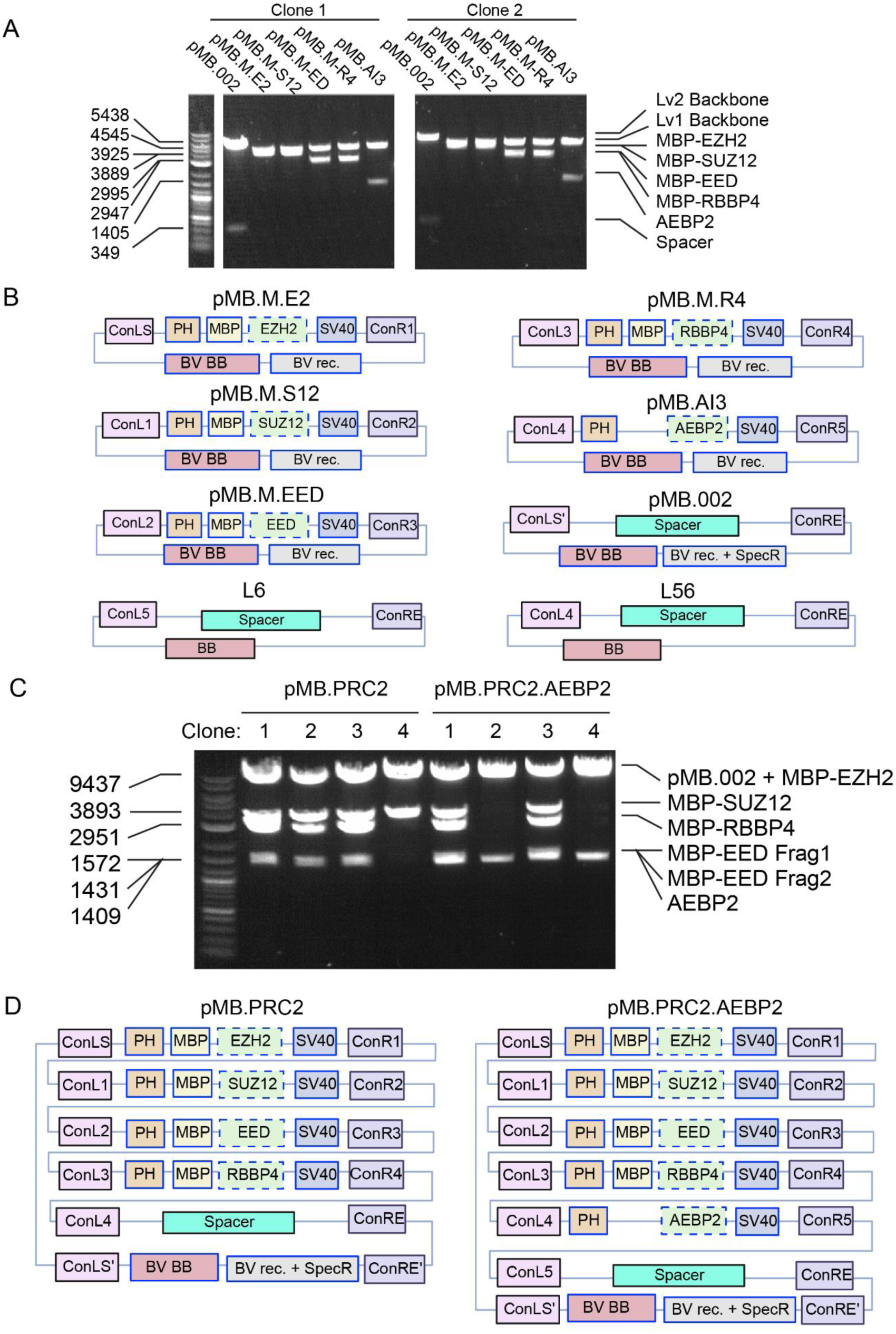
Proof-of-principle: construction of multigene baculovirus expression vectors for the expression of 4- and 5-subunit PRC2 complexes. (A) Golden Gate assembly of Level 1 plasmids was carried out using BsaI and two of the resulting clones were proceeded for restriction digestion using BsmBI. The plasmids pMB.M.E2, pMB.M.S12, pMB.M.ED, pMB.M.R4 and pMB.AI3 are Level 1 baculovirus expression vectors for the expression of EZH2, SUZ12, EED, RBBP4 and AEBP2 (isoform 3). The plasmid pMB.002 is a Level 1 plasmid that was assembled to contribute to the backbone during subsequent assemblies of Level 2 plasmids. (B) Schematic representation of the plasmids resulted from the Golden Gate assembly reaction that was analysed in (A). Presented are also two ‘linker plasmids’, L6 and L56, that are Level 1 plasmids that include a Spacer and Connector parts from the MoClo Yeast toolkit [18], designed to connect the last Transcription Unit to the backbone during the assembly of a Level 2 plasmid. (C) Golden Gate assembly using the Level 1 plasmids from (B) was carried out using BsmBI to construct Level 2 plasmids for a baculovirus expression of the 4-subunit PRC2 (left) or the 5-subunit PRC2-AEBP2 (right). Left: pMB.M.E2, pMB.M.S12, pMB.M.ED, pMB.M.R4, pMB.002 and L56 were assembled to form an expression vector coding for PRC2. In this context, the linker L56 was designed to connect the 4th transcription unit to the backbone. Right: PRC2-AEBP2 expression vector was assembled from the same Level 1 plasmids, with the exception that the Level 1 plasmid pMB.AI3 was added to include also the expression of AEBP2 and the L56 linker plasmid was replaced by the L6 linker plasmid, to connect the 5th transcription unit to the backbone. Four clones were selected from each Golden Gate reaction and were subjected to restriction digestion using MfeI and SalI. (D) Schematics of the obtained Level 2 multigene expression vectors, as indicated. In (B) and (D), boxes with a blue outline represent parts that originated from new Level 0 plasmids from the MoClo Baculo toolkit, while black outlined boxes are from the MoClo Yeast toolkit [18]. Dashed blue boxes represent the CDS parts generated in this study but are not a part of the MoClo Baculo toolkit. PH: polyhedrin promoter; SV40: SV40 polyA terminator; MBP: maltose-binding protein with PreScission cleavage site; Spacer: DNA sequence that was designed to fill up a gap during vector building (originated from the MoClo Yeast toolkit [18]); BV BB, BV rec, and BB are the same as in Fig. 1; BV rec + SpecR: Tn7 recombination sites compatible with the Bac-to-Bac baculovirus expression system, in the presence of Spectinomycin resistance. The Level 1 plasmids presented in (B) confer Ampicillin and Gentamicin resistance in *E. coli*. The Level 2 plasmids presented in (D) confer Ampicillin, Gentamicin and Spectinomycin in *E. coli*. In (A) and (B), fragments were analysed on 0.7% agarose TAE gel.

As with other systems of Modular Cloning [16,18], the efficiency of assembling Level 2 vectors may be reduced with the size and the number of parts. Here we demonstrate the construction of two Level 2 vectors, for the baculovirus expression of the PRC2 core complex in the presence or absence of its accessory subunit AEBP2 (Fig. 2C,D). Specifically, a 19 kbp PRC2 expression vector was designed for the expression of a 4-protein complex (MBP-EZH2, MBP-SUZ12, MBP-EED and MBP-RBBP4) and assembled with 75 % efficiency (3/4 colonies; Fig. 2C). A 21 kbp PRC2-AEBP2 expression vector designed to express five protein subunits (MBP-EZH2, MBP-SUZ12, MBP-EED, MBP-RBBP4 and AEBP2) and was assembled with an efficiency of 50% (2/4 colonies; Fig. 2C).

Each of the Golden Gate assemblies that were carried out to generate the expression vectors for the 4- and 5-subunit PRC2 complexes included a linker plasmid (L6 and L56 in Fig 2B). Such a linker plasmid is required when less than 6 Transcription Units are incorporated and aims to connect the last Transcription Unit to the backbone. The linker plasmid is assembled in a Golden Gate reaction using BsaI, by combining an ampicillin resistant *E. coli* Backbone plasmid (BB in Fig. 1A and Supplementary Table 1; plasmid pYTK095 from the MoClo Yeast Toolkit), a Spacer (parts 2-4, plasmid pYTK048 from the MoClo Yeast toolkit) and the appropriate Connector parts (parts 1 and 5; Supplementary Table 1) that are required to connect the last Transcription Unit to the Level 2 Backbone plasmid (pMB.002; Supplementary Table 2).

The sizes of the two Level 2 plasmids that we generated—19 kbp and 21 kbp—approach the upper size of plasmids that can be efficiently assembled using Golden Gate assembly (commonly reported as 20-30 kbp [16,18,19]). The expression of all subunits is driven by the same polyhedrin promoter and SV40 terminator, similar to biGBac [14] and Multibac systems [12]. Selected Level 2 clones were subjected to whole plasmid sequencing before downstream protein expression and purification.

As a proof of principle for vector construction compatible with the biGBac system, we assembled a baculovirus vector aiming for the expression of an eight subunit complex (Supplementary Fig. 3). This was done first with the MoClo Baculo system, using Golden Gate assembly, to construct the three Level 2 vectors (Supplementary Fig. 3B): pMB.BIG1a.PRC2 (expresses four PRC2 subunits), pMB.BIG1b.MTF2-PALI1 (for the expression of two accessory subunits of PRC2: MTF2 and PALI1), and pMB.BIG1c.G9A-GLP (for the expression of the H3K9me2 methyltransferases G9A and GLP). The resulting three Level 2 vectors are compatible with the biGBac system, which was used to combine them using Gibson Assembly into one Level 3 vector of 32 kb (pBIG2abc.PRC2.1-EHMT), aiming for the co-expression of all eight subunits (Supplementary Fig. 3C). The efficiency of constructing the Level 3 vector was 33% (2 out of 6 clones were correct; Supplementary Fig. 3A). The produced Level 3 vectors were verified using whole plasmid sequencing. We found sporadic mutations in the Gibson Assembly homology regions. These mutations were likely introduced during the Gibson Assembly reaction and are not expected to affect protein expression. This is because the homology regions are nonfunctional in Level 3 plasmids, as at this stage the Gibson Assembly has already been completed. No other mutations were identified in these Level 3 plasmids, which were assembled correctly.

### Proof of principle: protein expression and purification

As proof of principle, we used the PRC2- and PRC2-AEBP2 multigene baculovirus expression vectors that were constructed above (pMB.PRC2 and pMB.PRC2.AEBP2, respectively; Fig. 2D) for protein expression and purification. The obtained macromolecular complexes were pure, assembled and monodispersed (Fig. 3A-D,G). We obtained about 4 mg and 1.5 mg of PRC2 and PRC2-AEBP2, respectively, per litre of Hi5 insect cell culture. We also demonstrated PRC2 protein expression from the biGBac-compatible vector pMB.BIG1a.PRC2 (Fig. 3H), which was assembled solely by Golden Gate assembly using MoClo Baculo (Supplementary Fig. 3B).

**Figure 3.**
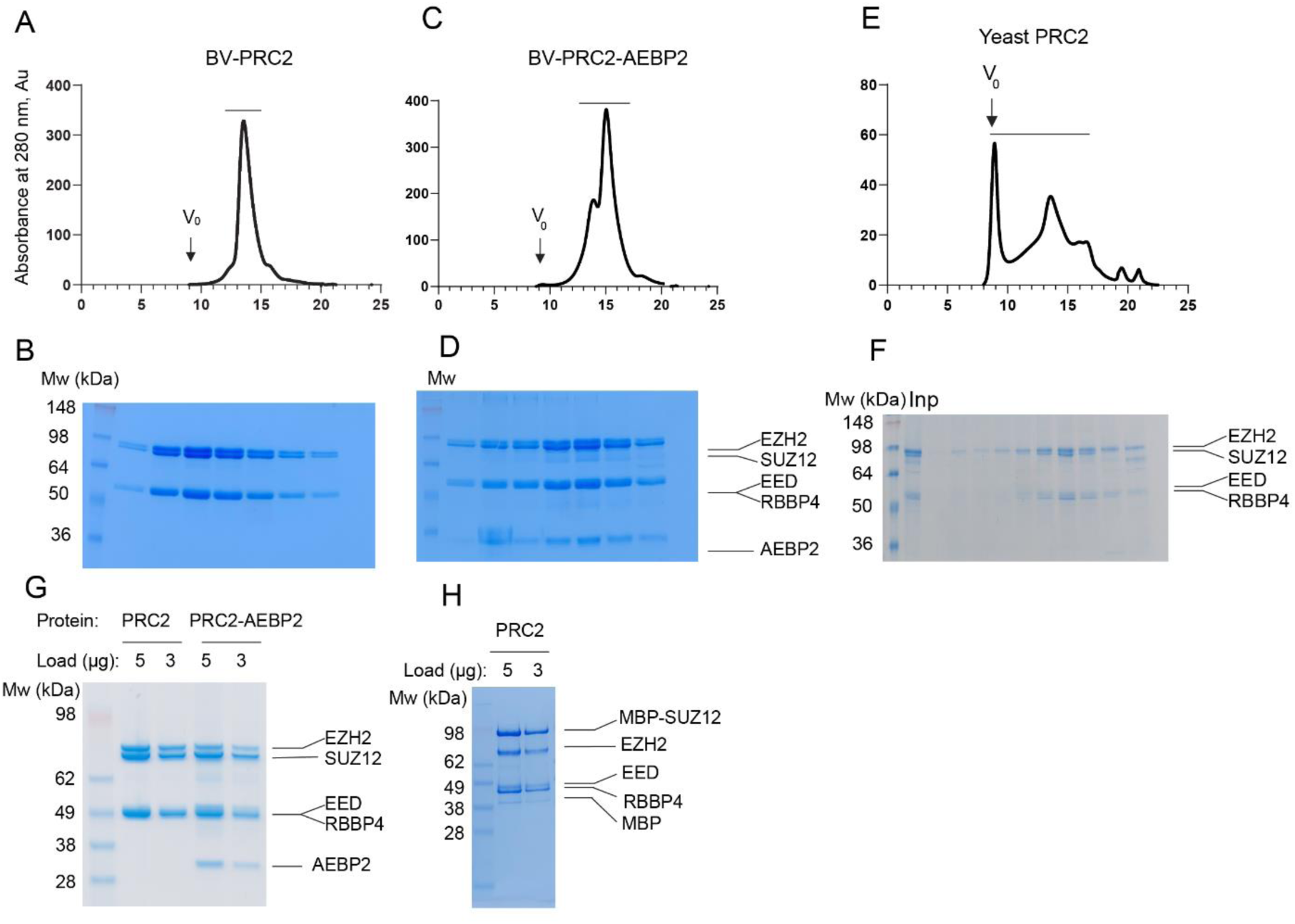
MoClo Baculo-expressed 4- and 5-subunit PRC2 complexes are pure and assembled. (A) PRC2 was expressed from baculovirus (BV) using the pMB.PRC2 multi-gene expression vector (Fig. 2D, left) and was fractionated over a Superose 6 Increase 10/300 column. UV absorbance at 280 nm was plotted to trace protein elution. The void volume is marked as V_0_. The black horizontal line above the peak indicates fractions that were taken for SDS-PAGE analysis in (B). (B) PRC2 gel filtration fractions from (A) were analysed by SDS-PAGE (Tris-Glycine, 12% acrylamide) and stained by coomassie. (C) PRC2-AEBP2 expressed using pMB.PRC2.AEBP2 construct (Fig. 2D, right) was analysed by Superose 6 Increase 10/300 column. (D) Fractions marked by the black line in (C) were analysed by SDS-PAGE (Tris-Glycine, 12% acrylamide). (E) A PRC2 yeast expression construct (pMY.PRC2; see Supplementary Fig. 4 for vector construction) was integrated into the genome of *S. cerevisiae* using homologous recombination for protein expression and then PRC2 that was purified and fractionated over a Superose 6 Increase 10/300 column. (F) Fractions that are marked by the horizontal black line in (E) were analysed on an SDS-PAGE (Tris-Glycine, 12% acrylamide). “Inp” indicates a sample before the size exclusion chromatography. (G) Purified PRC2 and PRC2-AEBP2 from (A) and (C), respectively, were analysed on SDS-PAGE (Bis-Tris, 4%-12% acrylamide) after fractions were pooled and proteins were concentrated. (H) Amylose affinity purification of PRC2 from lysate of SF9 cells that were infected with baculovirus carry the pMB.BIG1a.PRC2 vector (Supplementary Fig. 3B), analysed on SDS-PAGE (Bis-Tris, 8%-16% acrylamide).

As a proof-of-principle of protein expression using the yeast *S. cerevisiae*, we generated an expression vector for PRC2 (pMY.PRC2; Supplementary Fig. 4). For vector building, we used the same CDS (part 3) and Connectors (parts 1 and 5) parts that were used to build the baculovirus expression vectors above. To allow for yeast expression, we replaced the promoters (part 2/2a) and terminators (part 4) with *S. cerevisiae* promoters and terminators. For the same reason, we replaced the parts responsible for integration into the host DNA (parts 4a-d) with parts that were designed for integration into *S. cerevisiae* chromosome in yeast (Fig. 1A and Supplementary Fig. 4). We also replaced the MBP tags by a dual Protein A tag (Protein A) that we find more suitable in the yeast system, possibly given the lower expression level of human PRC2 in S. *cerevisiae* (more below). We then integrated the multi-gene PRC2 expression vector into *S. cerevisiae* chromosome through homologous recombination and used the obtained strain for the expression of the human PRC2. The obtained PRC2 was then purified as an assembled complex (Fig. 3E,F).

PRC2 is a histone methyltransferase enzyme. Therefore, we wished to determine that the recombinant PRC2 protein complex that we obtained is enzymatically active *in vitro*. To this end, we carried out an *in vitro* histone methyltransferase assay, to compare the enzymatic activities of the recombinant 4-subunits human PRC2 complexes that we purified using three different approaches: (i) Baculovirus expression, through the co-infection of four baculovirus stocks that were made using standard pFastBac1 vectors, each designed to express a single PRC2 subunit (Fig. 4, “pFB1 PRC2”), including the protein subunits MBP-EZH2, MBP-SUZ12, MBP-RBBP4 and MBP-EED. (ii) Baculovirus expression using a single MoClo Baculo-derived multi-gene vector carries Transcription Units for the same four PRC2 subunits with the same MBP tags (Fig. 4, “MoClo Baculo PRC2, insect cells”). (iii) A yeast expression of PRC2 using a single multi-gene expression vector including four Transcription Units of the same PRC2 subunits as above, with the exception that EZH2 carries a 2xProtein A tag and the rest of the subunits are untagged (Fig. 4, “MoClo Baculo PRC2, yeast”). In all cases, all tags were removed during the protein purification process and were not present in the enzymatic assay.

**Figure 4.**
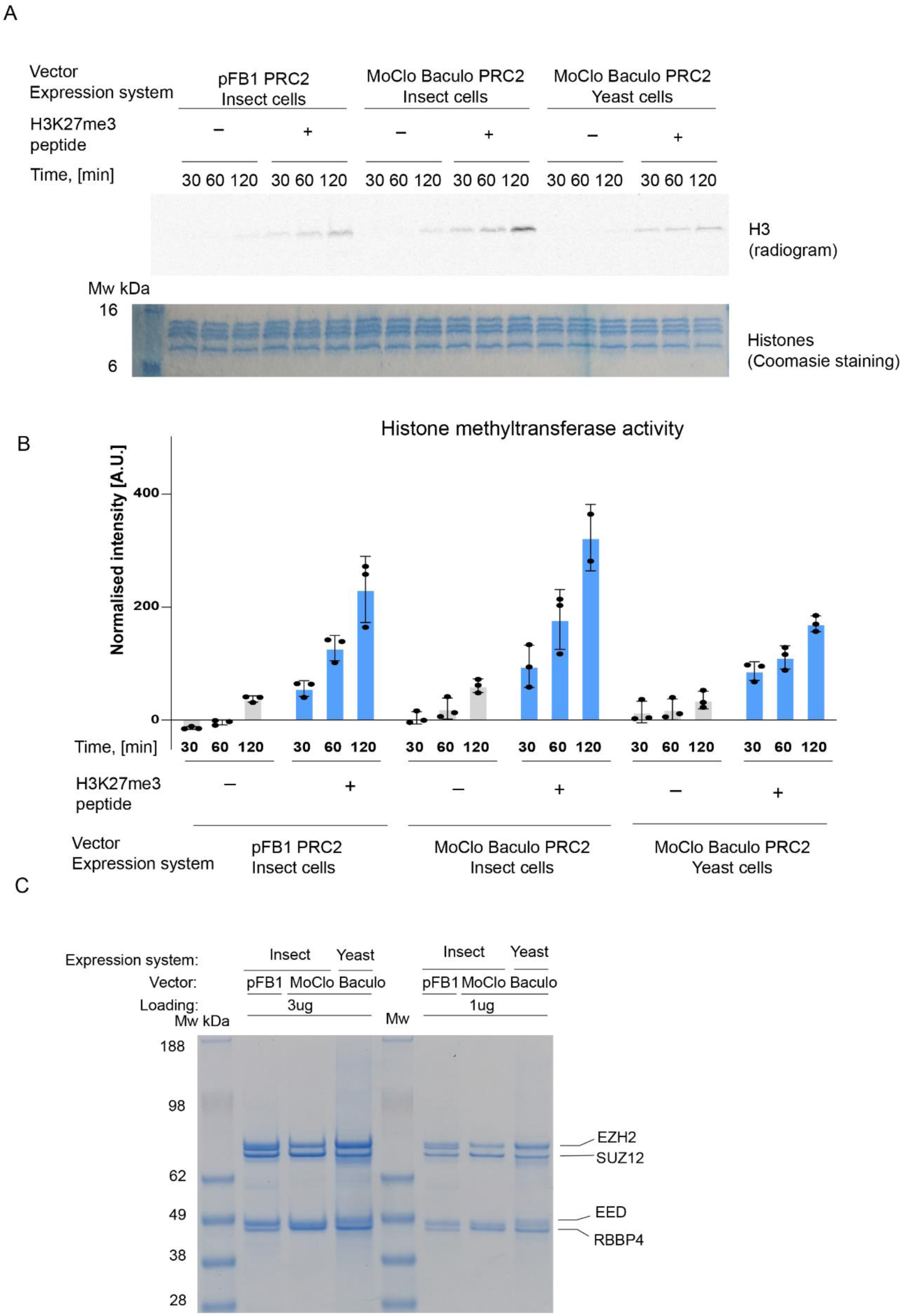
MoClo Baculo-expression vectors designed either for the baculovirus system or for yeast produced recombinant human PRC2 complexes that are enzymatically active. A. HMTase assay performed with 600 nM PRC2, 600 nM mononucleosomes, 5 µM S-Adenosyl-L-methionine [methyl-^14^C] and in the presence or absence of 50 µM H3K27me3 peptide, as indicated. The reaction mixture was separated with SDS-PAGE, stained with coomassie and exposed to a phosphor screen to produce a radiogram. (B) The bar plot (bottom) represents the relative HMTase activities of PRC2 in the absence (grey) or presence of H3K27me3 (blue) peptides, as indicated. Individual observations are presented, mean represents the normalised densitometry and error bars represent standard deviation of three independent experiments that were carried out on different days. (C) Equal amounts (1 or 3 μg) of the PRC2 complexes that were used for the HMTase assays were analysed on an SDS-PAGE (Tris-Acetate, 3%-8% acrylamide). In all panels: “insect” and “yeast” referred to protein that was expressed using the baculovirus expression system or *S. cerevisiae*, respectively. “pFB1’’ referred to a PRC2 complex that was expressed by the co-infection of 4 baculovirus stocks, each made of one pFastBac1 plasmid coding for one protein. “MoClo Baculo’’ referred to PRC2 complexes that were expressed from either baculovirus or yeast expression vectors, as indicated, constructed using parts of the MoClo Baculo toolkit.

All the PRC2 samples exhibited a substantial enzymatic activity only in the presence of an allosteric effector peptide H3K27me3 (Fig. 4B, blue bars), in agreement with PRC2 being an allosteric enzyme [20]. We also observed the automethylation of the catalytic subunit EZH2 (Supplementary Fig. 5A,B), which has to be methylated for PRC2 to become active [21,22].

We observed some variations in the enzymatic activity of the different recombinant human PRC2 preparations. These variations in activity were rather small, within <2-fold of the apparent reaction rate (Fig. 4B). These variations in activity might be attributed to the different expression systems. We also cannot exclude slight variations in the quantification of the protein concentration or subunit stoichiometry between samples, although SDS-PAGE did not identify large differences between them (Fig 4C). Overall, we demonstrated the construction of multi-gene expression vectors using MoClo Baculo and their subsequent application of protein expression and purification using the baculovirus system and yeast.

## Discussion

The MoClo Baculo system allows a high flexibility in building expression vectors while altering tags and expression systems as needed. According to data herein (Table 1) and elsewhere [23, 24], the baculovirus system provides a higher protein yield per litre than yeast. On the other hand, from the point plasmids are constructed, protein expression using the baculovirus system is a slow process (20-30 days under ideal conditions), while the yeast expression system provides a faster progress (6-7 days; Table 1 and [24]). Another advantage of the yeast expression system is the lower cost of the media: roughly 40-20% of the per-litre cost compared to the baculovirus system (Table 1). We propose that one could use MoClo Baculo in order to construct expression vectors for fast and cost-effective protein expression trials in yeast. Subsequently, the best-performing set of ORFs and tags can be assembled into baculovirus expression vectors, aiming for a high-yield protein production in insect cells.

**Table 1.** Comparison between the Baculovirus system to yeast in terms of protein expression. Values represent the best estimate based on the expression of the 4-subunit PRC2 complex from multigene expression vectors generated using MoClo Baculo. ^a^ ‘Protein expression’ and ‘Expression cost’ are defined here as the process from the stage the plasmid is made until the cells that expressed the protein are pelleted and collected, but before the protein purification process has started. For the baculovirus system, this process includes the preparation, amplification and quantification of the baculovirus particles, insect cell culture buildup, infection, incubation during the protein expression period and cell harvest. For the yeast system, this process includes transformation and colony growth, media prep, culture starter and inoculation, induction, growth and cell harvest. Expression trial in yeast was carried out using an *S. cerevisiae* line that was generated through the homologous recombination of the multi-gene expression vector into a chromosome. Estimated costs are provided in US dollars. ^b^ In this work, protein in *S. cerevisiae* was carried out using a galactose-inducible promoter on one of the PRC2 subunits (EZH2). This strategy added a substantial cost because of the usage of lactate (expression suppressor) and galactose (expression inducer). A usage of constitutive promoter, when feasible, would avoid galactose and lactate and therefore reduce the expression cost by about 50%. ^c^ Labour was estimated in the form of cumulative working hours that were dedicated to the process of protein expression, as defined in ^a^.

|  | **Baculovirus** | **Yeast** |
| --- | --- | --- |
| Time required to build a multi-gene (Level 2) plasmid | 5 days | 5 days |
| Protein expression <sup>a</sup> | 20-30 days | 6-7 days |
| Yield per litre (PRC2) | 3-5 mg | 0.03-0.05 mg |
| Cost of protein expression <sup>a</sup> | \$ 17/mg | \$ 500-830/mg |
| Expression cost per batch <sup>a</sup> | \$ 270 per 4 L batch | \$ 300 per 12 L batch <sup>b</sup> |
| Estimated labour per batch <sup>c</sup> | 4.6 h per 4 L batch | 4.8 h per 12 L batch |

A benefit of MoClo Baculo, with respect to most other baculovirus systems (Table 2), is that the entire vector construction process can be automated, allowing high throughput cloning. Specifically, MoClo Baculo is PCR free, which reduces the likelihood of mutations and eliminates the need for primer design and PCR optimization. Open reading frames can be synthesised as plasmids or gene fragments (e.g. gBlocks), such that they can be directly domesticated into Level 0 vectors by Golden Gate assembly. For the same reason, MoClo Baculo would be beneficial also by reducing the time and training required to design and construct complex expression vectors in any lab.

**Table 2.** Comparison between MoClo Baculo to other baculovirus expression systems. *The construction of up to Level 2 plasmids with up to 6 transcription units is done using Golden Gate assembly, even if the Level 2 plasmids are compatible with the biGBac system. Yet, the optional connection of multiple biGBac-compatible Level 2 plasmids with up to 6 transcription units each into a single Level 3 plasmids of up to 30 transcription units is done using the biGBac system and required Gibson Assembly. **Pre-build parts support up to 6 ORFs per MoClo Baculo vector (Level 2) and up to 30 ORFs if biGBac compatible parts are used to connect multiple Level 2 vectors into biGBac vectors (Level 3 here). Yet, the number of ORFs per MoClo Baculo vectors (Level 2), and therefore the number of ORFs in Level 3 vectors, can be expended if more MoClo Baculo Connector parts are designed, as in the case of the MoClo Yeast Toolkit [18]. However, we find that the Golden Gate assembly of Level 2 plasmids becomes very inefficient when the final product size approaches 30 kbp.

|  | **MoClo Baculo** | **Multibac [12]** | **Bac-to-Bac [6]** | **BiGBac [14]** | **GoldenBac [15]** |
| --- | --- | --- | --- | --- | --- |
| **Expression system** | BV and Yeast (*S. cerevisiae*) | BV | BV | BV | BV |
| **Modular Cloning** | Yes | No | No | Modular for the multigene level, but not for building the transcription units. | Modular for the multigene level, but not for building the transcription units. |
| **PCR or Gibson Assembly required** | N* | Y | Y | Y | Y |
| **Maximal number of ORFs per expression vector** | 6 (or 30 through compatibility with biGBac)** | Theoretically unlimited | 2 | 25 | 15 |

As for other MoClo cloning standards, MoClo Baculo would allow labs and institutes to manufacture, store and share collections of ORFs and tags ready for their assembly into multi-gene expression vectors for baculovirus and yeast. MoClo Baculo was designed as a general cloning standard, with the end goal of expression and purification of multi-subunit complexes for applications of structural biology, biophysics and biochemistry. As such, MoClo Baculo could potentially be expanded to other protein expression systems, as done with other MoClo tools for various applications [25,26,27,28,29]. Overall, MoClo Baculo is a cost-effective, fast and robust system for constructing multi-gene expression vectors, demonstrated here using the baculovirus system and yeast.

## Acknowledgements

We thank the support of the Micromon Genomics facility. Z.L. is supported by the Monash Graduate Scholarship, Monash International Tuition Scholarship. This work was supported by Sylvia and Charles Viertel Senior Medical Research Fellowship (C.D.) and the National Health and Medical Research Council (NHMRC) grant numbers APP1162921, APP1184637, APP2011767 and APP2020900 (C.D.).

## Author contributions

Z.L., M.B. and C.D. designed the project, Z.L., S.F.F. and M.B. carried out experiments while Z.L. led aspects related to the baculovirus system and M.B. led aspects related to yeast, Z.L. and M.B. analyzed data, Z.L., M.B. and C.D. wrote the manuscript and C.D. supervised the project.

## Methods

### Plasmid sequences

Part plasmids (Level 0) and expression vectors (Level 1 and 2) that were generated in this study are listed in Supplementary Table 2 and their annotated sequences are provided in GenBank file format in Supplementary Data 1. Plasmid sequence annotation was carried out with the aid of pLannotate [30]. Detailed information about plasmid construction is provided below. Even if one wishes to use MoClo Baculo exclusively for protein expression using the baculovirus system they would still need some plasmids from the MoClo Yeast toolkit, such as the Level 0 backbone (plasmid pYTK001), Connector part plasmids, Spacer part (plasmid pYTK048) and an *E. coli* backbone (plasmid pYTK095). The MoClo Yeast toolkit plasmids are available on AddGene (Kit #1000000061) and their sequences are provided in [18].

### Design of coding sequences (CDS) for MoClo Baculo

All CDS sequences were designed for compatibility with the MoClo Yeast system [18]. This includes the addition of overhangs that are reserved for part 3 (CDS; Supplementary Fig. 1C). Flanking BsmBI and BsaI sites were added to the CDS sequence, designed to generate the desired overhands for Golden Gate assembly into Level 0 and Level 1 plasmids, respectively (See Supplementary Fig. 1D).

In the MoClo Baclo system, part 3 allows for protein expression either with or without an N-terminal tag (Supplementary Fig. 1A), so only one CDS part is needed per project. As with the MoClo Yeast toolkit, the design of the CDS involved the introduction of synonymous mutations to remove the restriction sites for NotI, BsmBI, and BsaI from within the protein open reading frame (i.e., with the exception of the flanking BsmBI and BsaI sites). If one wishes to use MoClo Baculo to construct biGBac plasmids, they will also have to ensure that CDS and tag parts do not include PmeI sites, although this would rarely be a problem as PmeI sites are extremely uncommon. If one wishes only to construct expression vectors for the baculovirus system, not for yeast, then it is not required to remove NotI sites from CDS parts. This is because NotI is only used to linearise plasmids before their homologous recombination into a yeast chromosome. Finally, CDS are then domesticated into Level 0 plasmids, as described in the next section.

### Generation of new part (Level 0) plasmids

This process does not need to be repeated in cases where part plasmids already exist. It would typically be carried out once when a new part (e.g. a new tag) is introduced for the first time. The appropriate part-specific flanking sequences (Supplementary Fig. 1C, Supplementary Table 3) and universal flanking sequences (Supplementary Fig. 1D, Supplementary Table 3) were added to part sequences, either through gene synthesis or PCR primers, to enable domestication into the Level 0 vector and the subsequent modular cloning. DNA with the sequence of the promoter, terminator and tag parts were obtained either as ampicillin-resistant plasmids from gene synthesis (GenScript), gene fragments (IDT gBlocks), or PCR amplicons (Phusion® High-Fidelity DNA Polymerase, NEB) that were generated from plasmids with a pFastBac1.HMBP backbone. The parts were then cloned using Golden Gate assembly (BsmBI-v2, NEB #R0739) into the Level 0 backbone plasmid (pYTK001) of the MoClo yeast toolkit. The Backbone part (BV BB, Part 6b-c for the baculovirus system; Supplementary Table 1; plasmid pMB.001) for Level 1 MoClo Baculo baculovirus expression plasmids, was synthesised by Genscript to remove the BsaI and BsmBI sites and insertion of GFP-dropout.

### Generation of level 2 backbone plasmids

pMB.002 was generated by the golden gate assembly with BsaI enzyme using pMB.001 and pMB.004 from the MoClo Baculo toolkit, together with pYTK048, pYTK008, pYTK073 from yeast MoClo toolkit [18]. To construct the biGBac compatible backbones pMB.BIG1a to pMB.BIG1e (Supplementary Fig. 2A), the same reaction was performed using modified pYTK008 plasmids (ConLS’A to ConLS’E, respectively) and modified pYTK073 plasmids (ConLE’B to ConLE’F, respectively), where the biGBac compatible homology regions for Gibson Assembly (sequences A to F [14]; Supplementary Table 4) were inserted by BsmBI linearisation and HiFi DNA assembly (NEB, #E2621L) using ssDNA templates.

### Golden Gate assembly using BsaI and BsmBI

For Golden Gate assembly using BsaI, 25 fmol of each plasmid was added per reaction and assembled with NEBridge® Ligase Master Mix (NEB #M1100) and BasI-HF-V2 (NEB #R3733) according to protocol provided by the manufacturer. For Golden Gate assembly using BsmBI, 25 fmol of each plasmid was added per reaction with 1 µLBsmBI-v2 (NEB #R0739), 1 µL HiT4 ligase (NEB #R2622), 1.5 µL 10x T4 ligase buffer (NEB #B0202), 1 µL 500 mM NaCl making up a 15 µL reaction. The mixture was cycled 60 times between 16 °C and 50 °C for 5 min each, followed by 30 min final ligation at 16 °C and digestion at 50 °C for 15 min. The reaction was held at 10 °C and stored at -20 °C until use.

XL1-Blue (Agilent) was used as the only bacterial strain for all the cloning. Transformation and culturing of XL1-Blue is carried out according to the manufacturer’s protocol. For the propagation of pMB.001, the transformed cells were selected on an ampicillin plate. For the selection of Level 1 clones, we find that it is easier to identify GFP positive colonies (i.e. where Golden Gate assembly failed) on ampicillin plates.

### Construction of Level 3 plasmids using the biGBac system (optional)

pBIG2 plasmids were linearised in PmeI (NEB #R0560) restriction digest reaction at a final concentration of 10 fmol/µL. All multigene-carrying pMB.BIG1a-e plasmids were miniprepped (Qiagen #27106) with an extra PE buffer wash and eluted in 40 µL 60 °C Milli-Q water twice. Then, 500 fmol of each plasmids were mixed in a total of a 10 µL reaction of 1x Cutsmart (NEB) and with 1 µL PmeI (NEB #R0560). If plasmid concentration was not sufficient, all plasmids were mixed and dried in speedvac and re-dissolved in 9 µL 1x Cutsmart (NEB). All digestions were carried out at 37 °C for 90 minutes. Then, 1 µL of linearised backbone and 1 µL digested pMB.BIG1a-e mix were used to set up a 20 µL Gibson Assembly (NEB #E2611) reaction. The ligation mix were used to transform chemical-competent NEB10-beta (NEB #C3019) or electro-competent XL1-Blue, depending on the size of the final product. Generally, for plasmid between 20-30 kbp, electroporation is beneficial but not essential. For plasmid >30 kbp, electroporation is required.

### Expression and purification of PRC2 using the baculovirus system

Baculovirus was generated using Sf9 cells according to the instructions by the manufacturer (ThermoFisher). The virus titer was determined by the viability of infected SF9 cells using the MTS kit (Promega #G3580).

Protein expression was carried out according to the method previously described by Zhang et al [11]. Specifically, High Five cells were grown in Insect-XPRESS (Lonza #12-730Q) media and were infected with baculovirus at a cell density of 1.8-2.0*10^6 cells/ml. 70 hours post-infection, cell pellets were harvested at 1500 g and lysed in lysis buffer (50 mM Tris pH8 at 25 °C, 300 mM NaCl, 15 % glycerol, 0.5 % NP-40, 1 mM TCEP, cOmplete protease inhibitor (Roche #11836170001)) for 30 min at 4 °C. The lysate was clarified by centrifugation at 30,000 g for 30 min and the supernatant was incubated with amylose resin (NEB #E8021) for 60 min with rotation. Then, the resin was washed with 5 column volumes (c.v.) of lysis buffer, 10 c.v. of HSW buffer (50 mM Tris pH8 at 25 °C, 500 mM NaCl) and 10 c.v. of LSW buffer (50 mM Tris pH8 at 25 °C, 150 mM NaCl). The protein was eluted from the resin with LSW buffer supplemented with 10 mM maltose and 1 mM TCEP. Then the protein was digested overnight with PreScission protease and loaded to a 5 ml prepacked Heparin column (Cytiva #17040701) and eluted with a linear gradient over 10 c.v., starting from Buffer A (150 mM NaCl, 20 mM Tris pH 8.0 at 25 °C) to 50% Buffer B (2000 mM NaCl, 20 mM Tris pH 8.0 at 25°C). Finally, the eluate was diluted 1:1 with 20 mM Tris pH 8.0, 1 mM TCEP, concentrated using 30 kDa Amicon Ultra centrifugal filter and injected on to a Superose 6 Increase 10/300 column (Cytiva, #29091596) in HEPES pH7.5 at 25 °C and 150 mM NaCl. Fractions that contained protein of interest were supplemented with 1 mM TCEP, concentrated and frozen in liquid nitrogen. The entire purification was carried out at 4-8 °C.

### Expression and purification of PRC2 using S. cerevisiae

For the purification of the 4-subunits PRC2 from yeast, cells were picked from a single colony and grown overnight in -Ura synthetic minimal media (2 g/L Yeast Synthetic Drop-out Medium Supplement without uracil (Merck #1501), 67 g/L Yeast Nitrogen Base Without Amino Acids (Merck #Y0626)) supplemented with 3% glucose. The following day, cells were washed twice and used to inoculate 12 L of YP-lactate (1.5% Yeast extract (Merck #Y1625), 3% Meat Peptone (ThermoScientific H26694.36), 2% Lactate (Chem Supply #LL008)) in 6 flasks. Cultures were grown at 30 °C and 180 rpm for 31 h. EZH2 expression was induced by the addition of 3% galactose and the cultures were grown for 19 h. Cells were harvested, washed in water, pelleted (335 g), flash-frozen and stored at -80 °C until the protein purification was carried out. For protein purification, the cell pellet was thawed in an equal volume of 2X lysis buffer (400 mM NaCl, 50 mM Tris pH 7.5, 0.5 mM DTT, 17 mg/mL phenylmethanesulfonyl fluoride (PMSF; Merck #P7626), 25 μg/mL leupeptin (Merck #2884), 125 μg/mL pepstatin A (Merck #516481), 33 mg/mL benzamidine (Merck #B6506)), and lysed using a DynoMill. Lysate was clarified by centrifugation at 30,000 g, followed by ultracentrifugation at 45,000 rpm. The supernatant was bound overnight to 20 mL of IgG resin. Beads were washed with a high salt buffer (500 mM NaCl, 20 mM Tris pH 7.5, 0.5 mM DTT), followed by a low salt buffer (200 mM NaCl, 20 mM Tris pH 7.5, 0.5 mM DTT). PRC2 complex was eluted using an overnight cleavage by TEV protease (100 μg) and subjected to heparin purification. The heparin peak was concentrated and injected into a Superose 6 10/300 column (Cytiva, #29091596). The protein in the peak corresponding to PRC2 was concentrated and flash-frozen.

### SDS-PAGE

To resolve all subunits using SDS-PAGE, 3-8% Tris-Acetate gel (ThermoFisher #EA0375BOX) and 1X MES SDS Running buffer (ThermoFisher #NP0002) were used. 3 µg or 1 µg PRC2 complexes were supplemented to a final concentration of 1X LDS sample buffer (ThermoFisher #NP0007) with 1% 2-mercaptoethanol (Sigma-Aldrich #M3148) and heated at 95 °C for 5 min before loading onto a Tris-Acetate gel. The gel was run for 60 min at 160 V before staining with InstantBlue Coomassie protein stain (Expedeon, #ISB1L).

### Nucleosome reconstitution

Histones, octamers, and polynucleosome reconstitution were carried out as previously described [11]. In brief, recombinant histones were purified from inclusion bodies and reconstituted into histone octamers. 182 bp DNA was amplified using Pfu DNA polymerase and purified by ion exchange chromatography and isopropanol precipitation. Chromatin was assembled using gradient salt dialysis at 4°C. Chromatin was reconstituted by initially titrating across a range of octamer ratios (from 1:0.9 to 1:1.1 DNA:octamer molar ratio) in a 20 µL mixture of 3 µM DNA, 20 mM Tris pH 7.5 at 25 °C, 2 M NaCl, 1 mM EDTA and 1 mM DTT. For each of these samples, gradient salt dialysis was used at 4°C in a dialysis device (ThermoFisher #69572), starting from refolding buffer (20 mM Tris pH 7.5 at 25 °C, 2 M KCl, 1 mM EDTA and 1 mM DTT) to a medium salt buffer containing 20 mM Tris pH 7.5 at 25 °C, 250 mM KCl, 1 mM EDTA and 1 mM DTT over 18 h, and the final step dialysis was carried out using a low salt buffer containing 20 mM Tris pH 7.5 at 25 °C, 2.5 mM KCl, 1 mM DTT. The quality of mononucleosomes was assessed by 0.8 % agarose TBE and 6 % polyacrylamide TBE gel electrophoresis, and the most appropriate molar ratio of DNA:octamer was selected for a large-scale batch. Large-scale reconstitution was conducted as above, except in a volume of 0.5 mL using dialysis tubing (Spectrum #888-11527). The folded mononucleosome was concentrated with 500 µL 100 kDa MWCO concentrator (Cytiva, #28932237). The concentration of the mononucleosome was measured using a Qubit protein assay kit (ThermoFisher #Q3321). Mononucleosomes were stored at 4 °C until use.

### *In vitro* HMTase activity assays using radiolabeled S-adenosyl-L-methionine

For the HMTase reactions, each 10 µL reaction contained 0.6 µM PRC2 produced from either yeast or insect cells, 0.6 µM chromatinised gene, and 5 µM S-[methyl-14C]-adenosyl-L-methionine (PerkinElmer, #NEC363050UC). Reactions were incubated in reaction buffer (50 mM Tris-HCl pH 8.0 at 30°C, 100 mM KCl, 2.5 mM MgCl_2_, 0.1 mM ZnCl_2_, 2 mM 2-mercaptoethanol, 0.1 mg/mL BSA, and 5% v/v glycerol) for 30, 60, or 120 min at 30°C. Reactions were stopped by adding 4X LDS loading dye (Thermo Fisher Scientific, #NP0007) supplemented with 4% 2-mercaptoethanol (Sigma-Aldrich #M3148) to a final concentration of 1X LDS and heating at 95°C for 5 min. The reactions were then loaded onto 16.5 % SDS-PAGE gels and ran on ice for 120 min at 160 V in 1X Tris-glycine buffer. Gels were stained with InstantBlue Coomassie protein stain (Expedeon, #ISB1L) before vacuum-drying for 1 h at 60°C. Dried gels were then exposed to a storage phosphor screen for six days before acquiring radiograms using a Typhoon 5 Imager (GE Healthcare). All experiments were performed in three independent replicates that were carried out on three separate days. Densitometry was carried out using ImageLab software (Bio-Rad). Values were interpolated from a standard curve using GraphPad Prism 10.2.1 and the background was subtracted, then values plotted with SD displayed.

---

**Supplementary Figure 1.**
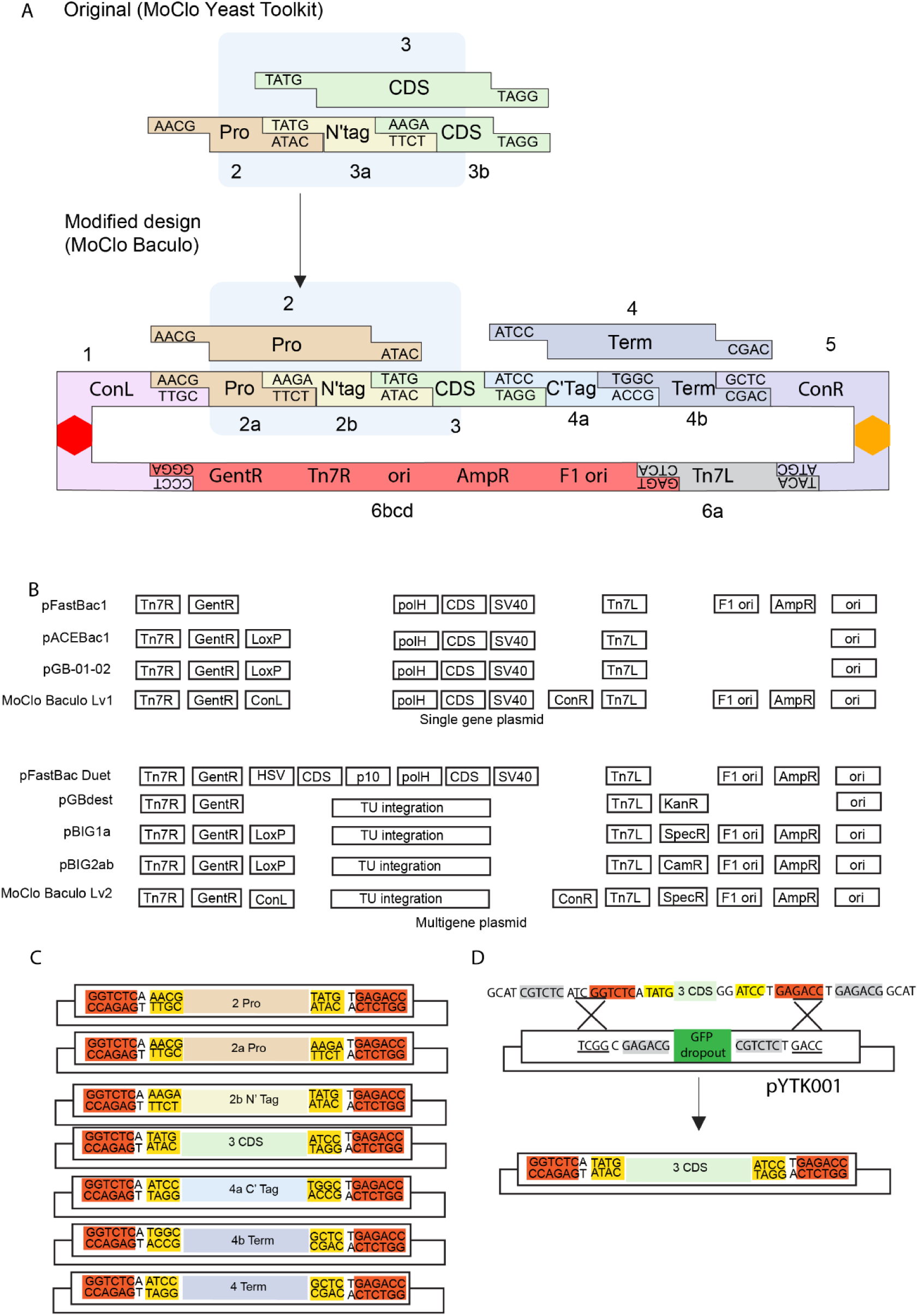
Level 1 plasmid architecture of the MoClo Baculo toolkit. (A) The MoClo Baculo Level 1 plasmid design is illustrated, including the left connector (ConL), promoter (Pro), N-terminal tag (N’tag), coding sequence (CDS), C-terminal tag (C’Tag), terminator (Term) and the right connector (ConR). Changes that were made to the original MoClo Yeast toolkit design [18] in order to enable the usage of the same CDS part either in the presence or absence of an N-terminal tag are indicated above (highlighted in blue boxes). Presented are also the four base pair overhangs that are generated in each part after BsaI digestion. The parts are numbered according to the MoClo Yeast numbering system, with the exception of parts that were modified (in the blue box) and the backbone parts that uses the same overhangs as in the MoClo Yeast toolkit but are all collectively numbered as part 6 for simplicity (specifically, the MoClo Baculo parts 6a, 6b, 6c and 6d were originally termed parts 6, 7, 8a and 8b, respectively, in the Moclo Yeast toolkit, while in the MoClo Baculo toolkit part 6bcd is a single part that include all the baculovirus plasmid backbone elements, as indicated, with the exception of the Tn7L element that is included in part 6a). Parts 2-4 are related to the expressed gene, part 6 allows recombination into baculovirus DNA or into the yeast genome, and parts 1 and 5 are for the integration into multi-gene vectors (Level 2 plasmids). The two colourful hexagons represent nucleotide sequences designed to adhere to complementary sequences in other connectors, after the subsequent digestion of the plasmid with type IIS restriction enzymes (see Fig. 1A-B). (B) A comparison between the organisation of different functional modules of various baculovirus expression vectors that are compatible with the Bac-to-Bac expression system for the expression of single (top) or multiple (bottom) genes and the single-(Level 1; “Lv1”) and multi-gene (Level 2; “Lv2”) MoClo Baculo expression vectors. Presented are the Tn7 left and right recombination sites (Tn7R and Tn7L), gentamicin, kanamycin, spectinomycin, ampicillin and chloramphenicol resistance (GenR, KanR, SpecR, AmpR and CamR), LoxP recombination site (LoxP), left and right connectors for Golden Gate assembly (ConL and ConR), a region that hosts multiple genes or transcription units (TU integration), Simian virus polyadenylation signal (SV40), Herpes Simplex Virus thymidine kinase polyadenylation signal (HSV), coding sequence (CDS), baculovirus p10 and polyhedrin promoters (p10 and polH), f1 origin (F1 ori), and pUC origin (ori). (C) Flanking sequence of promoter (Pro), N-terminal tag (N Tag), coding sequence (CDS), C-terminal tag (C’Tag), terminator (Term) within the context of Level 0 vectors. BsaI restriction site is highlighted in orange and the overhang in yellow. Overhang sequences are inherited from the MoClo yeast toolkit, with the exception of parts 2a, 2b and 3, which were redesigned. (D) In order to generate Level 0 plasmids that holds a given part (presented in panel (C)), the part can be obtained as a double strand DNA (e.g. as a gene fragment or a plasmid from gene synthesis) while including flanking sequences suitable for its Golden Gate Assembly into the pYTK001 plasmid (originated from the MoClo Yeast Tool Kit [18]) using BsmBI. As an example, presented are the complete flanking sequences that are added to a CDS part (“3 CDS” representing an open reading frame sequence) that is ordered from gene synthesis. BsmBI restriction sites are highlighted in grey, the overhang for assembly with pYTK001 is underlined and the rest of the colours are as in panel (C). The black crosses represent the site of ligation during BsmBI golden gate reaction.

**Supplementary Figure 2.**
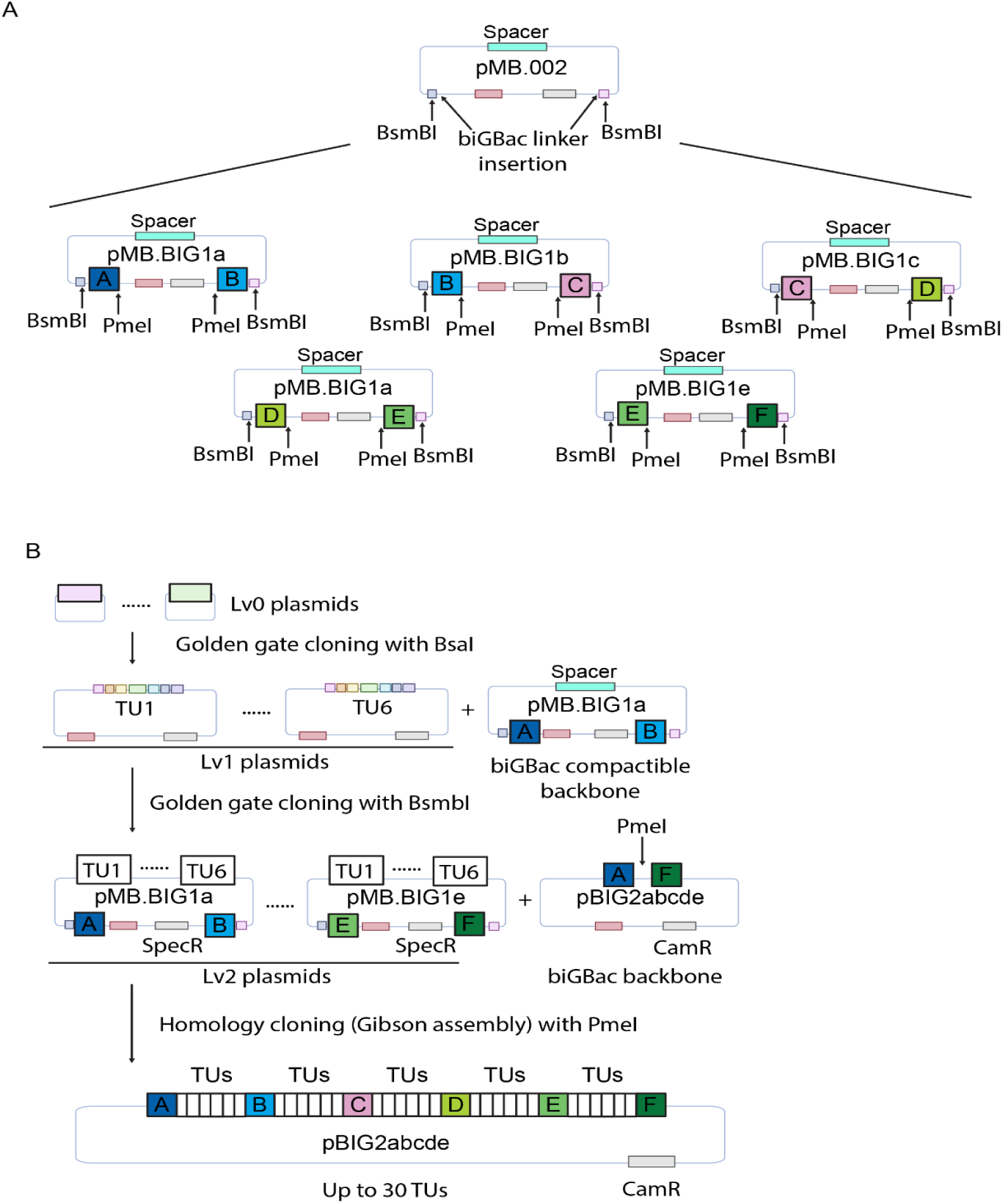
MoClo Baculo level 2 plasmids can be connected into larger Level 3 plasmids through compatibility with the biGBac system. (A) pMB.BIG1a to pMB.BIG1e plasmids (bottom) were generated by inserting biGBac linker sequence (box marked with A to F) and PmeI site between the pMB.002 plasmid backbone and BsmBI site. (B) The workflow of constructing biGBac compatible level 2 vectors is essentially the same as other MoClo Baculo vectors, except that the pMB.BIG1a to pMB.BIG1e plasmids are used as Level 2 backbone instead of the pMB.002 plasmid. Colour key is as in Fig. 1 with addition of biGBac linker sequences A to F in assorted colours. TU: transcriptional unit; TUs: BsmBI golden gate ligated transcriptional units; SpecR: spectinomycin resistance; CamR: chloramphenicol resistance.

**Supplementary Figure 3.**
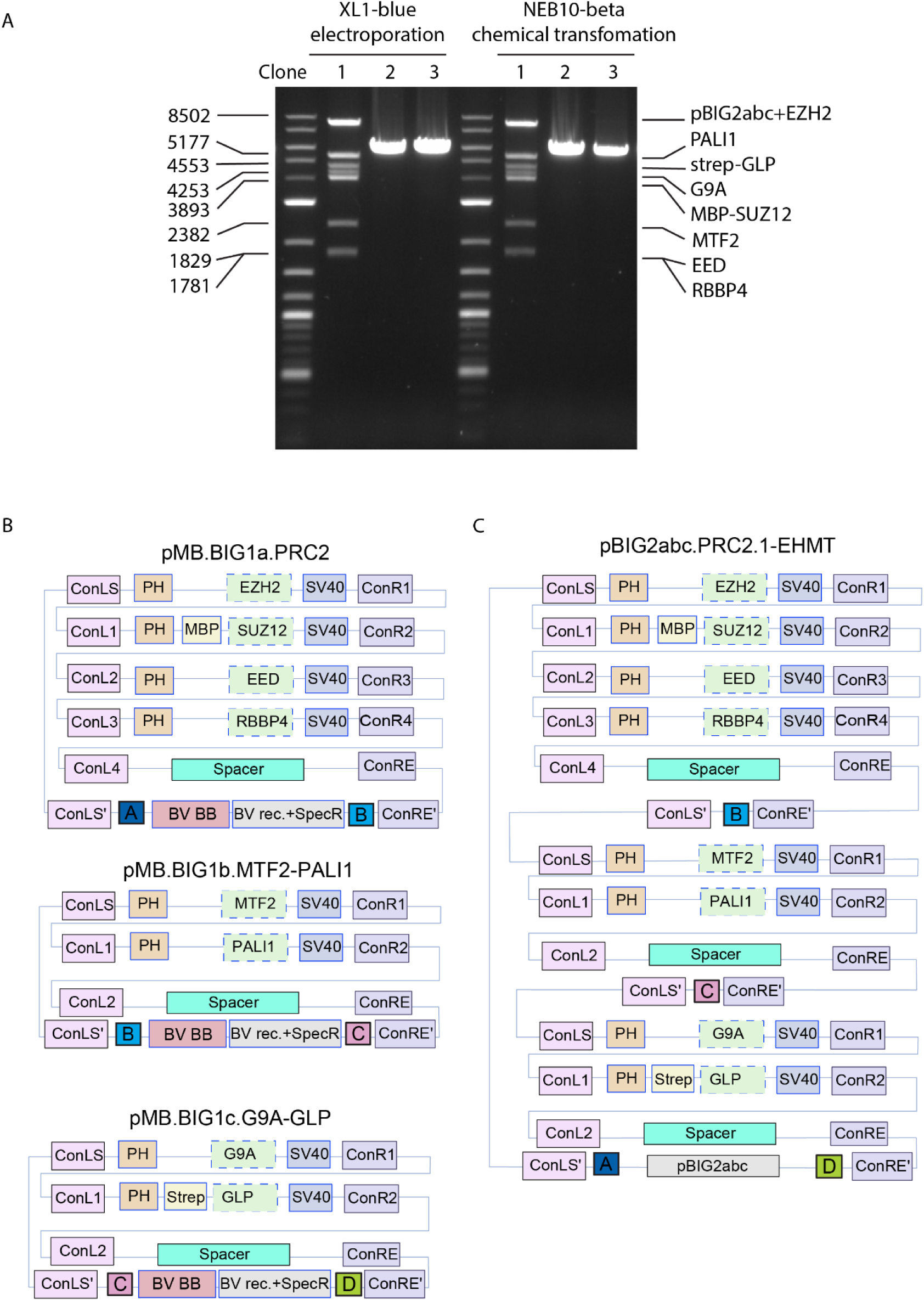
Proof-of-principle: Construction of a Level 3 vector through compatibility between the MoClo Baculo to the biGBac system. **(A)** Gibson assembly of pMB.BIG1a.PRC2, pMB.BIG1b.MTF2-PALI1 and pMB.BIG1c.G9A-GLP plasmids were carried out using pBIG2abc as backbone. The final plasmid, pBIG2abc.PRC2.1-EHMT, is validated by restriction digestion using MfeI and PmeI. Chemical- and electroporation transformation were benchmarked here. Clone 1 are correct clones, while 2 and 3 are empty pBIG2abc. (B) Schematic representation of the plasmids used for the Gibson assembly reaction that was analysed in (A). (C) Schematic representation of the plasmids produced by the Gibson assembly reaction was analysed in (A).

**Supplementary Figure 4.**
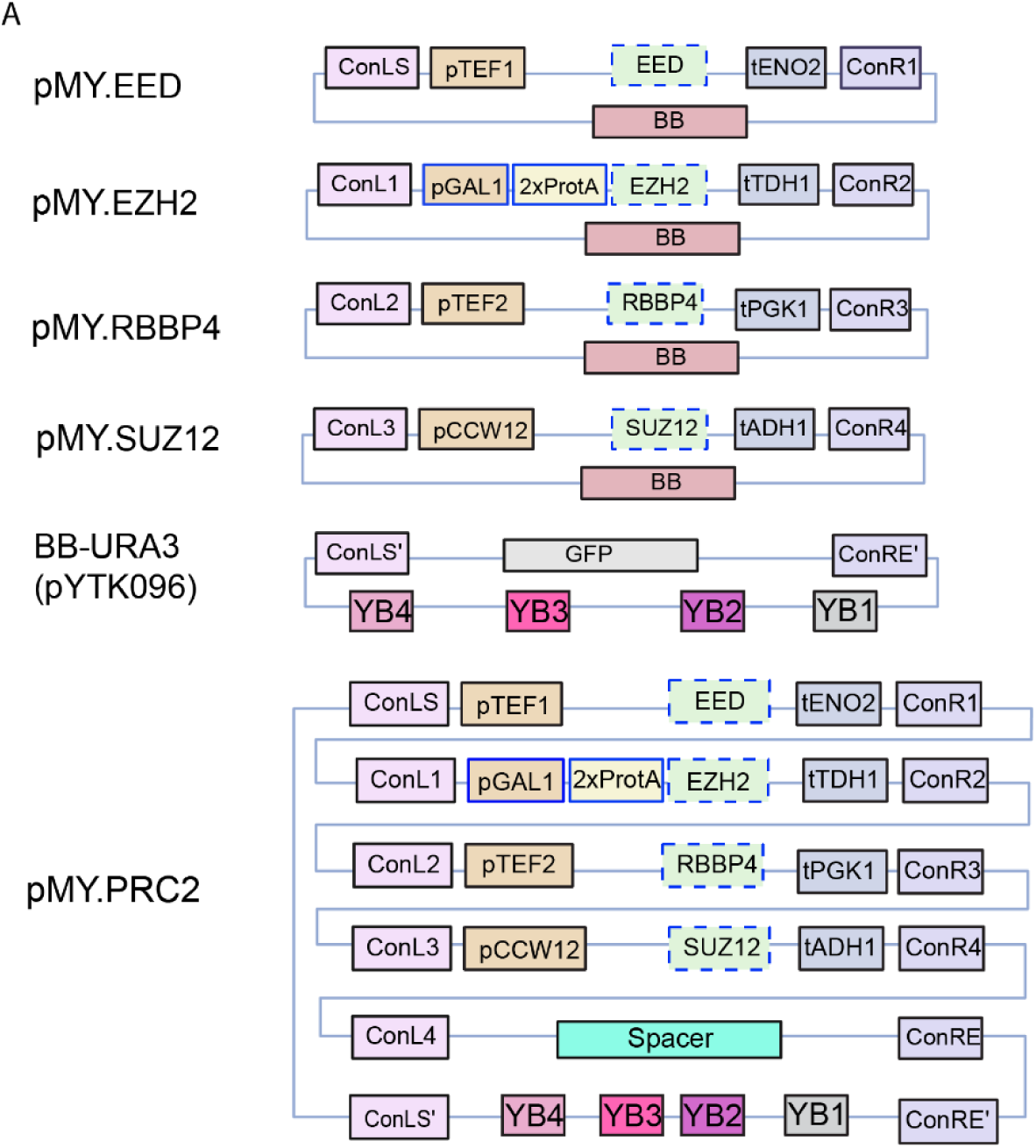
A construct design of a yeast expression plasmid. Schematic of the Level 1 and Level 2 constructs that were made using parts of the MoClo Baculo and MoClo Yeast toolkits [18] for the expression of the human PRC2 in *S. cerevisiae*. Parts from the MoClo Yeast toolkit are in black boxes and parts of the MoClo Baculo (parts 2b and 2a; see in Supplementary Fig. 1A) are in blue boxes. Dashed blue boxes represent the same CDS parts that were also used to construct baculovirus expression vectors in Fig. 2. “BB-URA3” mark a backbone plasmid that originated from the MoClo Yeast toolkit [18] (pYTK096), aiming to enable genomic integration into the *URA3* locus of the *S. cerevisiae* chromosome, using selection by uracil prototrophy. YB1: URA3 selection mark, YB2: URA3 3’Hom, YB3: Backbone (ori, KanR), YB4: URA3 5’Hom.

**Supplementary Figure 5.**
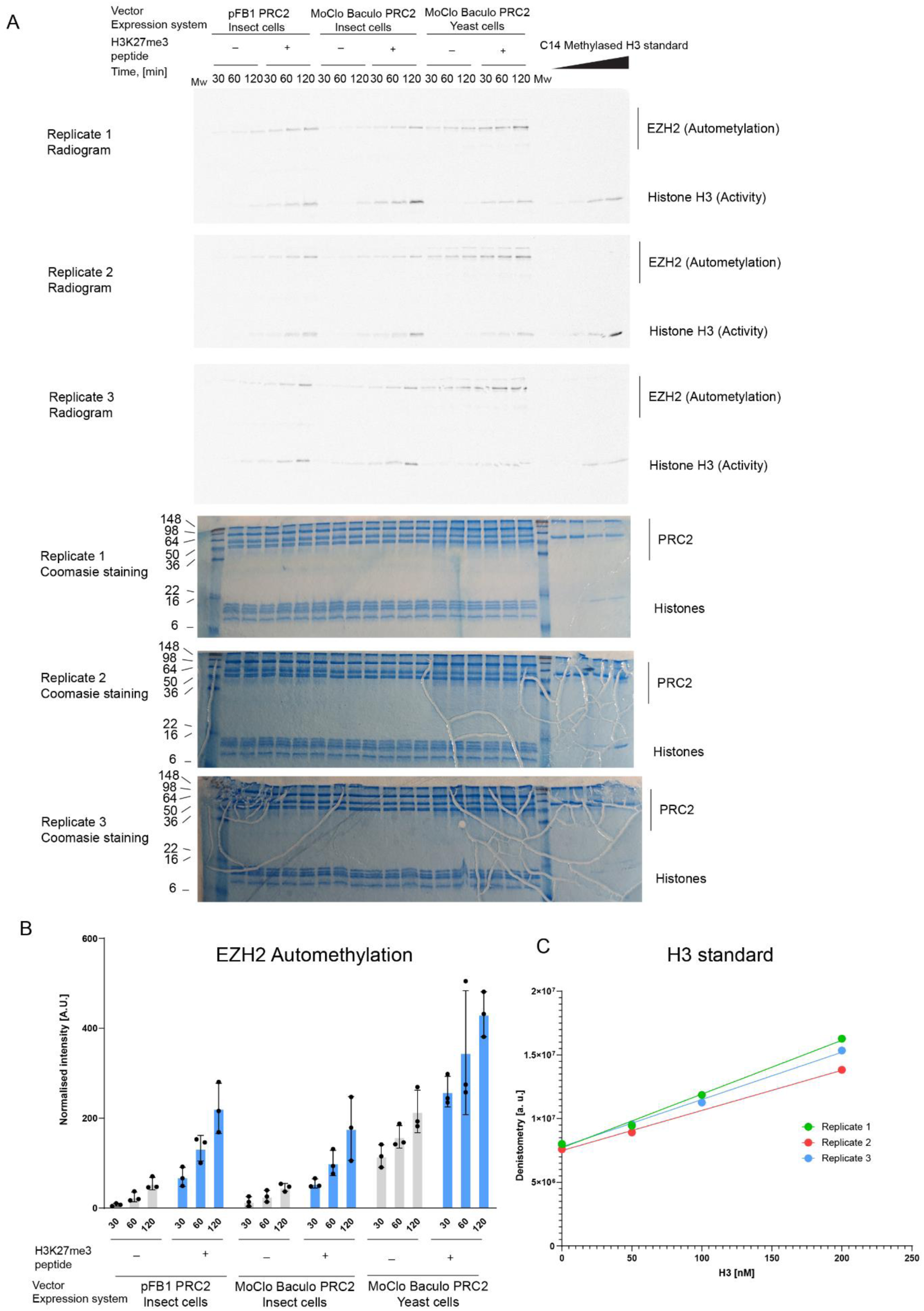
Human PRC2 complexes expressed using MoClo Baculo vectors in insect cells and in yeast are enzymatically active. (A) Uncropped radiogram and coomassie stained gels from three independent replicates of HMTase activity assay that were carried out on three different days (the cropped gels are in Fig. 4A). The ^14^C-labelled H3 standard that is used for normalisation between different replicates is shown on the right side of the gel. B. The EZH2 automethylation signal was quantified using ^14^C-labelled H3 standards and the values were plotted as a bar chart. C. H3 standard curves that were used to normalise data from independent replicates of HMTase activity assays. The standard curves of the different replicates are presented, with colour key as indicated.

## Legends for Supplementary Tables and Data

**Supplementary Table 1. Compatibility between parts for the construction of expression vectors for baculovirus and yeast.**

Tabulated are relevant parts (level 0 plasmids) from the MoClo Baculo and MoClo yeast toolkits, where the latter are indicated by an asterisk. Part numbers and sub-part numbers are as in Supplementary Fig. 1A. Expression system-specific parts, either for the baculovirus (BV) system or yeast, are clustered together (see left column). All the tags, coding sequences (CDS) and connector parts of the MoClo Baculo system are compatible with vector building for both yeast and baculovirus expression. Promoter, terminator and backbone parts are system-specific. In brackets, the names of type 6 parts as they appear in Fig. 1 and across the text.

**Supplementary Table 2. Plasmids that were generated during this study.** Indicated are plasmids that were constructed during this study, including new parts (Level 0 plasmids) and expression vectors (Level 1 and 2 plasmids). Excluded are vectors that were constructed as intermediate products of multi-gene vector buildings for the yeast expression system, as they were not designed to express proteins.

**Supplementary Table 3. Flanking sequences that are added when generating new parts.** As mentioned in the Methods section, new parts can be generated by the addition of BsmBI and BsaI sites that generate the correct overhang sequences upon cleavage. In this Table, indicated are specific sequences that need to be added upstream (“Upstream integration sequence”) or downstream (“Downstream integration sequence”) to a certain part sequence in order to form BsmBI and BsaI sites with the correct overhang. “Requirements” indicate functional elements that have to be added together with the part sequence to enable its functionality, as well as additional instructions for part design. Underlined are ATG start codons. Part 6bcd does not include a BsmBI site as the plasmid itself is a Level 0.

**Supplementary Table 4. Sequences of ssDNAs that were used to generate linker sequences for compatibility with the biGBac system.** Indicated are the sequences that were added to generate the modified ConLS’A to ConLS’E and ConRE’B to ConRE’F parts that were used as precursors for generating the biGBac compatible plasmids pMB.BIG1a to pMB.BIG1e, respectively.

**Supplementary Data 1. Plasmid sequences.** Sequences of baculovirus-specific Level 0 plasmids that were generated during this study are included within a zipped directory as GenBank files.

**Supplementary Data 2. Plasmid sequences.** Sequences of yeast-specific Level 0 plasmids that were generated during this study are included within a zipped directory as GenBank files.

**Supplementary Data 3. Plasmid sequences.** Sequences of Level 0 plasmids that are used for both yeast and the baculovirus system and that were generated during this study are included within a zipped directory as GenBank files.

**Supplementary Data 4. Plasmid sequences.** Sequences of ORFs as Level 0 plasmids that were synthesised during this study are included within a zipped directory as GenBank files.

**Supplementary Data 5. Plasmid sequences.** Sequences of single-gene and multi-gene baculovirus expression vectors (Level 1 and Level 2) plasmids, respectively, that were generated during this study are included within a zipped directory as GenBank files.

**Supplementary Data 6. Plasmid sequences.** The sequence of a yeast expression multigene plasmid (Level 2) that was generated during this study is included within a zipped directory as GenBank files.

## Reference

1. Saibil HR. Cryo-EM in molecular and cellular biology. Mol Cell. 2022 Jan 20;82(2):274–284. doi: 10.1016/j.molcel.2021.12.016. PMID: 35063096.

2. Baser B, van den Heuvel J. Assembling Multi-subunit Complexes Using Mammalian Expression. Adv Exp Med Biol. 2016;896:225–38. doi: 10.1007/978-3-319-27216-0_15. PMID: 27165329.

3. Assenberg R, Wan PT, Geisse S, Mayr LM. Advances in recombinant protein expression for use in pharmaceutical research. Curr Opin Struct Biol. 2013 Jun;23(3):393–402. doi: 10.1016/j.sbi.2013.03.008. Epub 2013 Jun 1. PMID:23731801.

4. Kost TA, Condreay JP, Jarvis DL. Baculovirus as versatile vectors for protein expression in insect and mammalian cells. Nat Biotechnol. 2005 May;23(5):567–75. doi: 10.1038/nbt1095. PMID: 15877075; PMCID: PMC3610534.

5. Baghban R, Farajnia S, Rajabibazl M, Ghasemi Y, Mafi A, Hoseinpoor R, Rahbarnia L, Aria M. Yeast Expression Systems: Overview and Recent Advances. Mol Biotechnol. 2019 May;61(5):365–384. doi: 10.1007/s12033-019-00164-8. PMID: 30805909.

6. Anderson D, Harris R, Polayes D. Rapid generation of recombinant baculoviruses and expression of foreign genes using the Bac-to-Bac baculovirus expression system. 1996; Focus 17:53–58

7. Sokolenko S, George S, Wagner A, Tuladhar A, Andrich JM, Aucoin MG. Co-expression vs. co-infection using baculovirus expression vectors in insect cell culture: Benefits and drawbacks. Biotechnol Adv. 2012 May-Jun;30(3):766-81. doi: 10.1016/j.biotechadv.2012.01.009. Epub 2012 Jan 28. PMID: 22297133; PMCID: PMC7132753.

8. Aucoin MG, Perrier M, Kamen AA. Improving AAV vector yield in insect cells by modulating the temperature after infection. Biotechnol Bioeng. 2007 Aug 15;97(6):1501–9. doi: 10.1002/bit.21364. PMID: 17274066.

9. Aucoin MG, Perrier M, Kamen AA. Production of adeno-associated viral vectors in insect cells using triple infection: optimization of baculovirus concentration ratios. Biotechnol Bioeng. 2006 Dec 20;95(6):1081–92. doi: 10.1002/bit.21069. PMID: 16952153.

10. Meghrous J, Aucoin MG, Jacob D, Chahal PS, Arcand N, Kamen AA. Production of recombinant adeno-associated viral vectors using a baculovirus/insect cell suspension culture system: from shake flasks to a 20-L bioreactor. Biotechnol Prog. 2005 Jan-Feb;21(1):154-60. doi: 10.1021/bp049802e. PMID: 15903253.

11. Zhang Q, McKenzie NJ, Warneford-Thomson R, Gail EH, Flanigan SF, Owen BM, Lauman R, Levina V, Garcia BA, Schittenhelm RB, Bonasio R, Davidovich C. RNA exploits an exposed regulatory site to inhibit the enzymatic activity of PRC2. Nat Struct Mol Biol. 2019 Mar;26(3):237–247. doi: 10.1038/s41594-019-0197-y. Epub 2019 Mar 4. PMID: 30833789; PMCID: PMC6736635.

12. Pelosse M, Crocker H, Gorda B, Lemaire P, Rauch J, Berger I. MultiBac: from protein complex structures to synthetic viral nanosystems. BMC Biol. 2017 Oct 30;15(1):99. doi: 10.1186/s12915-017-0447-6. PMID: 29084535; PMCID: PMC5661938.

13. Gibson DG, Young L, Chuang RY, Venter JC, Hutchison CA 3rd, Smith HO. Enzymatic assembly of DNA molecules up to several hundred kilobases. Nat Methods. 2009 May;6(5):343–5. doi: 10.1038/nmeth.1318. Epub 2009 Apr 12. PMID: 19363495.

14. Weissmann F, Petzold G, VanderLinden R, Huis In ‘t Veld PJ, Brown NG, Lampert F, Westermann S, Stark H, Schulman BA, Peters JM. biGBac enables rapid gene assembly for the expression of large multisubunit protein complexes. Proc Natl Acad Sci U S A. 2016 May 10;113(19):E2564-9. doi: 10.1073/pnas.1604935113. Epub 2016 Apr 25. PMID: 27114506; PMCID: PMC4868461.

15. Neuhold J, Radakovics K, Lehner A, Weissmann F, Garcia MQ, Romero MC, Berrow NS, Stolt-Bergner P. GoldenBac: a simple, highly efficient, and widely applicable system for construction of multi-gene expression vectors for use with the baculovirus expression vector system. BMC Biotechnol. 2020 May 12;20(1):26. doi: 10.1186/s12896-020-00616-z. PMID: 32398045; PMCID: PMC7216392.

16. Weber E, Engler C, Gruetzner R, Werner S, Marillonnet S. A modular cloning system for standardized assembly of multigene constructs. PLoS One. 2011 Feb 18;6(2):e16765. doi: 10.1371/journal.pone.0016765. PMID: 21364738; PMCID: PMC3041749.

17. Engler C, Gruetzner R, Kandzia R, Marillonnet S. Golden gate shuffling: a one-pot DNA shuffling method based on type IIs restriction enzymes. PLoS One. 2009;4(5):e5553. doi: 10.1371/journal.pone.0005553. Epub 2009 May 14. PMID: 19436741; PMCID: PMC2677662.

18. Lee ME, DeLoache WC, Cervantes B, Dueber JE. A Highly Characterized Yeast Toolkit for Modular, Multipart Assembly. ACS Synth Biol. 2015 Sep 18;4(9):975–86. doi: 10.1021/sb500366v. Epub 2015 May 1. PMID: 25871405.

19. Enghiad B, Xue P, Singh N, Boob AG, Shi C, Petrov VA, Liu R, Peri SS, Lane ST, Gaither ED, Zhao H. PlasmidMaker is a versatile, automated, and high throughput end-to-end platform for plasmid construction. Nat Commun. 2022 May 16;13(1):2697. doi: 10.1038/s41467-022-30355-y. PMID: 35577775; PMCID: PMC9110713.

20. Margueron R, Justin N, Ohno K, Sharpe ML, Son J, Drury WJ 3rd, Voigt P, Martin SR, Taylor WR, De Marco V, Pirrotta V, Reinberg D, Gamblin SJ. Role of the polycomb protein EED in the propagation of repressive histone marks. Nature. 2009 Oct 8;461(7265):762-7. doi: 10.1038/nature08398. Epub 2009 Sep 20. PMID: 19767730; PMCID: PMC3772642.

21. Lee CH, Yu JR, Granat J, Saldaña-Meyer R, Andrade J, LeRoy G, Jin Y, Lund P, Stafford JM, Garcia BA, Ueberheide B, Reinberg D. Automethylation of PRC2 promotes H3K27 methylation and is impaired in H3K27M pediatric glioma. Genes Dev. 2019 Oct 1;33(19-20):1428–1440. doi: 10.1101/gad.328773.119. Epub 2019 Sep 5. PMID: 31488577; PMCID: PMC6771381.

22. Wang X, Long Y, Paucek RD, Gooding AR, Lee T, Burdorf RM, Cech TR. Regulation of histone methylation by automethylation of PRC2. Genes Dev. 2019 Oct 1;33(19-20):1416–1427. doi: 10.1101/gad.328849.119. Epub 2019 Sep 5. PMID: 31488576; PMCID: PMC6771386.

23. Lowin T, Raab U, Schroeder J, Franssila R, Modrow S. Parvovirus B19 VP2-proteins produced in Saccharomyces cerevisiae: comparison with VP2-particles produced by baculovirus-derived vectors. J Vet Med B Infect Dis Vet Public Health. 2005 Sep-Oct;52(7-8):348-52. doi: 10.1111/j.1439-0450.2005.00871.x. PMID: 16316399.

24. Morton CL, Potter PM. Comparison of Escherichia coli, Saccharomyces cerevisiae, Pichia pastoris, Spodoptera frugiperda, and COS7 cells for recombinant gene expression. Application to a rabbit liver carboxylesterase. Mol Biotechnol. 2000 Nov;16(3):193-202. doi: 10.1385/MB:16:3:193. PMID: 11252804.

25. Moore SJ, Lai HE, Kelwick RJ, Chee SM, Bell DJ, Polizzi KM, Freemont PS. EcoFlex: A Multifunctional MoClo Kit for E. coli Synthetic Biology. ACS Synth Biol. 2016 Oct 21;5(10):1059–1069. doi: 10.1021/acssynbio.6b00031. Epub 2016 May 2. Erratum in: ACS Synth Biol. 2020 May 15;9(5):1225. doi: 10.1021/acssynbio.0c00177. PMID: 27096716.

26. Otto M, Skrekas C, Gossing M, Gustafsson J, Siewers V, David F. Expansion of the Yeast Modular Cloning Toolkit for CRISPR-Based Applications, Genomic Integrations and Combinatorial Libraries. ACS Synth Biol. 2021 Dec 17;10(12):3461–3474. doi: 10.1021/acssynbio.1c00408. Epub 2021 Dec 3. PMID: 34860007; PMCID: PMC8689691.

27. Iverson SV, Haddock TL, Beal J, Densmore DM. CIDAR MoClo: Improved MoClo Assembly Standard and New E. coli Part Library Enable Rapid Combinatorial Design for Synthetic and Traditional Biology. ACS Synth Biol. 2016 Jan 15;5(1):99–103. doi: 10.1021/acssynbio.5b00124. Epub 2015 Nov 4. PMID: 26479688.

28. Obst U, Lu TK, Sieber V. A Modular Toolkit for Generating Pichia pastoris Secretion Libraries. ACS Synth Biol. 2017 Jun 16;6(6):1016–1025. doi: 10.1021/acssynbio.6b00337. Epub 2017 Mar 15. PMID: 28252957.

29. Fonseca JP, Bonny AR, Kumar GR, Ng AH, Town J, Wu QC, Aslankoohi E, Chen SY, Dods G, Harrigan P, Osimiri LC, Kistler AL, El-Samad H. A Toolkit for Rapid Modular Construction of Biological Circuits in Mammalian Cells. ACS Synth Biol. 2019 Nov 15;8(11):2593–2606. doi: 10.1021/acssynbio.9b00322. Epub 2019 Nov 5. PMID: 31686495.

30. McGuffie MJ, Barrick JE. pLannotate: engineered plasmid annotation. Nucleic Acids Res. 2021 Jul 2;49(W1):W516–W522. doi: 10.1093/nar/gkab374. PMID: 34019636; PMCID: PMC8262757.

